# Paramyxovirus matrix proteins modulate host cell translation via exon-junction complex interactions in the cytoplasm

**DOI:** 10.1101/2024.09.05.611502

**Authors:** Chuan-Tien Hung, Griffin D Haas, Ruth E Watkinson, Hsin-Ping Chiu, Shreyas Kowdle, Christian S Stevens, Arnold Park, James A Wohlschlegel, Patricia A Thibault, Benhur Lee

## Abstract

Viruses have evolved myriad strategies to exploit the translation machinery of host cells to potentiate their replication. However, how paramyxovirus (PMVs) modulate cellular translation for their own benefit has not been systematically examined. Utilizing puromycylation labeling, overexpression of individual viral genes, and infection with wild-type virus versus its gene-deleted counterpart, we found that PMVs significantly inhibit host cells’ nascent peptide synthesis during infection, with the viral matrix being the primary contributor to this effect. Using the rNiV-NPL replicon system, we discovered that the viral matrix enhances viral protein translation without affecting viral mRNA transcription and suppresses host protein expression at the translational level. Polysome profile analysis revealed that the HPIV3 matrix promotes the association of viral mRNAs with ribosomes, thereby enhancing their translation efficiency during infection. Intriguingly, our NiV-Matrix interactome identified the core exon-junction complex (cEJC), critical for mRNA biogenesis, as a significant component that interacts with the paramyxoviral matrix predominantly in the cytoplasm. siRNA knockdown of eIF4AIII simulated the restriction of cellular functions by the viral matrix, leading to enhanced viral gene translation and a reduction in host protein synthesis. Moreover, siRNA depletion of cEJC resulted in a 2-3 log enhancement in infectious virus titer for various PMVs but not SARS-CoV-2, enterovirus D68, or influenza virus. Our findings characterize a host translational interference mechanism mediated by viral matrix and host cEJC interactions. We propose that the PMV matrix redirects ribosomes to translate viral mRNAs at the expense of host cell transcripts, enhancing viral replication, and thereby enhancing viral replication. These insights provide a deeper understanding of the molecular interactions between paramyxoviruses and host cells, highlighting potential targets for antiviral strategies.

## INTRODUCTION

Human parainfluenza virus type 3 is a member of the *Paramyxoviridae*^1,2^, a family of RNA viruses that includes pathogens of agricultural and global health importance such as Nipah virus (NiV), mumps virus, measles virus (MeV) and Newcastle disease virus (NDV)^1^. As a prominent respiratory tract pathogen, HPIV3 is a leading cause of various airway diseases, including pneumonia, croup, and bronchiolitis, with a notably high incidence in infants and young children ^3–7^. HPIV3 is characterized by a negative-stranded RNA genome encapsulated within a lipid envelope, which is derived from the host cell membrane. The genome encodes six principal genes: nucleocapsid (N), phosphoprotein (P), matrix (M), fusion (F), attachment (HN) proteins, and polymerase (L)^1,2^.

Paramyxoviruses (PMVs) are classic cytoplasmic replicating viruses, and the progeny virions are released from the plasma membrane of the host cell. Viral assembly and budding are orchestrated by the matrix protein (M), a major structural protein underlying the viral envelope^8–10^. Despite their cytoplasmic life cycle, paramyxoviral M proteins from diverse genera, including those from Nipah (NiV-M), Hendra (HeV-M), Ghana (GhV-M) and Cedar (CedV-M) viruses (genus *Henipavirus*), Sendai virus (SeV-M, genus *Respirovirus*), mumps virus (MuV-M, genus *Orthorubulavirus*) and Newcastle disease virus (NDV-M, genus *Orthoavulavirus*), exhibit nuclear-cytoplasmic trafficking that is essential for matrix function^11–22^. Notably, these proteins can be detected in the nucleus during the early stages of infection^16^. Furthermore, NiV-M and HPIV3-M have been shown to counteract antiviral Type I interferon through RIG- or mitophagy-mediated pathways, respectively^23–25^. These findings suggest that paramyxoviral M proteins may execute roles beyond viral assembly at the plasma membrane.

RNA viruses have adeptly evolved to exploit the machinery of host cells, a necessity stemming from their relatively limited genome capacity compared to DNA viruses. By co-opting the host cell machinery, they can potentiate their own replication. RNA viruses employ diverse strategies to inhibit host mRNA expression while selectively enhancing translation of viral mRNAs^26–28^. This targeted modulation of host and viral mRNA translation is believed to suppress antiviral responses, thereby facilitating viral replication within host cells. For instance, the SARS-CoV-2 Nsp1 binds to the ribosomal mRNA channel, inhibiting translation and inducing the degradation of translated cellular mRNAs, leading to a global reduction in protein translation^29–31^. Similarly, the Influenza A virus induces host translation shutoff by reducing the amount of host mRNA in cells^32–34^ and cap-snatching of cellular pre-mRNAs to prioritize viral transcripts^35,36^. Enteroviruses, such as poliovirus and coxsackievirus B3, use 2A proteinases to cleave eIF4G1 and PABP, resulting in rapid host translation shutoff and directing ribosomes to IRES-contained viral mRNAs^37–39^. Additionally, the matrix protein of the vesicular stomatitis virus (VSV) causes a global inhibition of host gene expression by interacting with the TFIIH transcription factor and forming a complex with Rae1 and Nup98 to disrupt mRNA export amongst other mechanisms yet to be defined^40–42^. In PMVs, parainfluenza virus 5 (PIV5) manipulates host cell translation through its P and V proteins, and the nuclear localization of matrix from NDV appears crucial for suppressing host cell transcription^43,44^. However, the specific mechanisms by which PMVs modulate host cellular translation for their own benefit remain obscure and have not been systematically examined.

Our previous studies have demonstrated the ubiquitin-regulated nuclear-cytoplasmic trafficking behavior of paramyxovirus matrix proteins, indicating a non-structural function within the nucleus that remains to be determined^14,16^. In this study, we explore an additional non-structural function of the viral matrix within the cytoplasm. We reveal that PMV matrix proteins inhibit host cell nascent peptide synthesis during infection by interacting with the core exon-junction complex (cEJC). This interaction enhances viral mRNA translation while suppressing host mRNA translation by promoting the association of viral mRNAs with ribosomes, thereby increasing their translation efficiency. The matrix-cEJC interaction occurs primarily in the cytoplasm, where matrix-induced re-localization of cEJC is observed during infection. Depletion of cEJC significantly enhances PMV replication, underscoring the critical role of this interaction. These findings suggest a model whereby PMV matrix redirects ribosomes to translate viral mRNAs, enhancing viral replication and providing insights into how HPIV3 manipulates host translation machinery.

## RESULTS

### HPIV3 matrix inhibits host protein synthesis during viral infection

To interrogate how paramyxovirus infection affects host protein synthesis, we utilized puromycylation to capture a snapshot of protein synthesis. The sensitivity of puromycylation in HEK-293Ts was first determined by treating cells with either mock or cycloheximide (CHX) to block translation, followed by a puromycin pulse to label newly synthesized proteins. CHX-treated cells showed >50% reduction in puromycin-incorporated proteins (Figure 1A, lane 3), demonstrating that puromycylation can quantitatively detect inhibited protein synthesis as expected^45^. We then evaluated the effect of HPIV3 infection on host protein synthesis. A 14% reduction in protein synthesis was noted at 24 hours post-infection (hpi) that became more pronounced at 48 hpi, approaching 35% decrease in synthesis compared to control cells (Figure 1B). An even more dramatic inhibitory effect on protein synthesis was evident in cells infected with Cedar virus (CedV) (Figure 1C, 82% decrease at 24 hpi), a non-pathogenic henipavirus within the *Paramyxoviridae* family. These results suggest that paramyxoviruses, such as HPIV3 and CedV employ strategies to disrupt host protein synthesis. Next, we sought to identify the viral determinants contributing to this disruption by examining cells expressing each FLAG-fused HPIV3 viral protein. Expression of the HPIV3 matrix led to a nearly 40% reduction in puromycylated proteins (Figure 1D, lane 4), whereas other viral proteins did not exhibit a similar effect. Notably, cells expressing nucleocapsid and phosphoprotein showed increased puromycylation, potentially reflecting their high expression levels. These findings suggest that the HPIV3 matrix plays a crucial role in inhibiting host protein synthesis.

**Figure 1.**
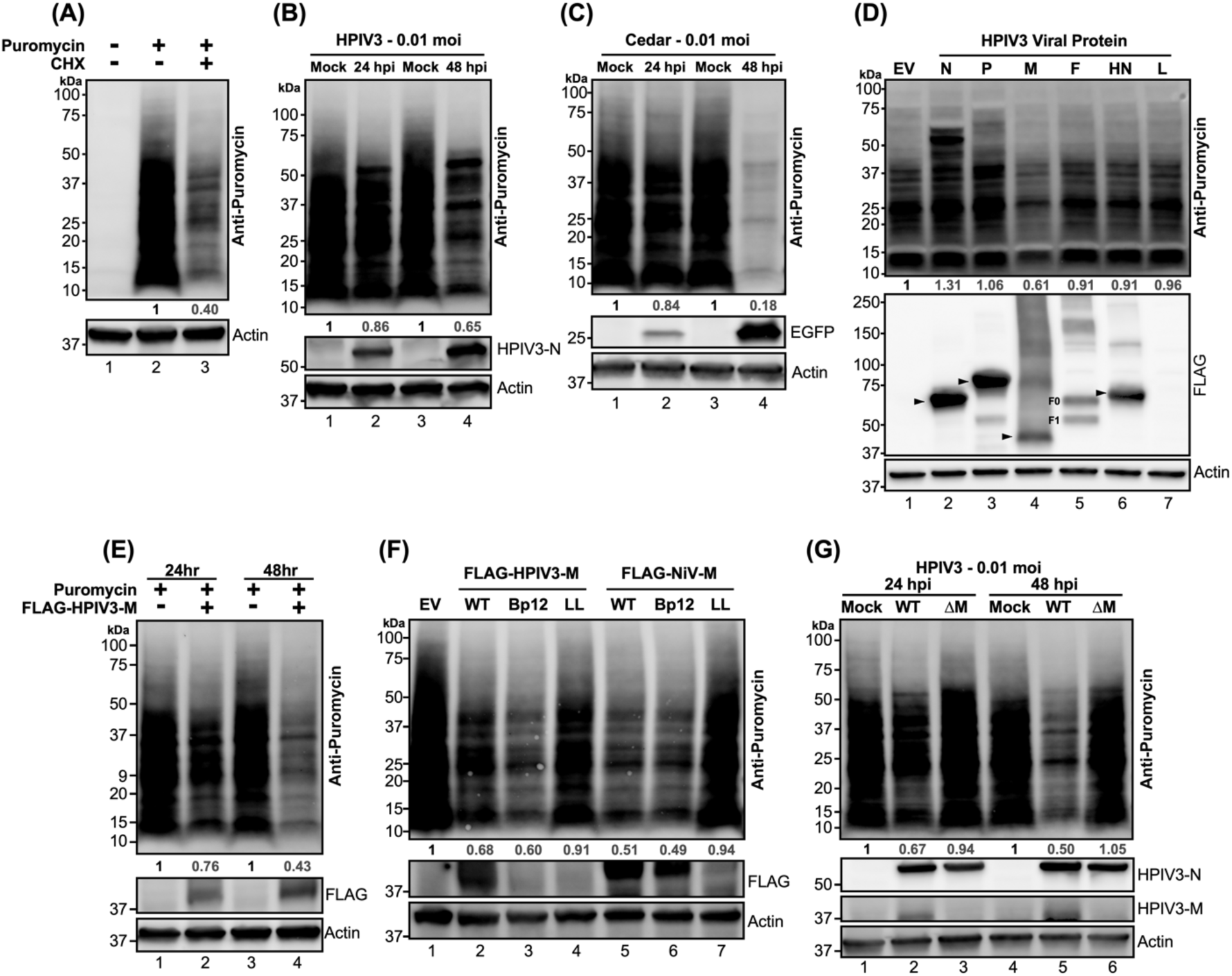
Puromycylation of newly synthesized protein in paramyxovirus-infected or viral proteins transfected HEK-293T cells. (A) HEK-293Ts were treated with either mock or cycloheximide (CHX) at 200 µg/ml for 5 hours, followed by a 20-minute treatment with either mock or puromycin at 10 µg/ml. After the puromycin pulse, cells were washed with PBS and re-fed with complete media. Lysates were analyzed by immunoblotting, and newly synthesized (puromycylated) proteins were probed using an anti-puromycin antibody. (B-C) HEK-293Ts were inoculated with mock, HPIV3, or Cedar virus and puromycin pulsed at indicated time points. Lysates were immunoblotted to probe puromycylated proteins. The expression of HPIV3 nucleocapsid (N) and EGFP served as controls for HPIV3 and Cedar infections, respectively. (D) Flag-fused viral proteins from individual HPIV3 genes or an empty vector (EV) were expressed in HEK-293T cells for 48 hours, followed by puromycylation and immunoblot analysis to detect puromycylated proteins. Expression of viral proteins was detected with an anti-FLAG antibody, with molecular weights indicated by black arrows. F0: F precursor. F1: cleaved F. (E-F) HEK-293T cells were expressed designated FLAG-fused matrix (M) proteins from HPIV3 or NiV, including wild-type (WT) and mutants (Bp12: NLS mutant, LL: NES mutant), along with EV control for 24 or 48 hrs. Following puromycylation, immunoblotting was conducted to determine the puromycylated protein and flag-fused matrix. (G) HPIV3 or HPIV3ΔM virus-infected HEK-293Ts were analyzed by immunoblotting after puromycylation at 24- and 48-hrs post-infection to detect puromycylated protein and HPIV3 viral proteins. HPIVP3-N and HPIV3-M served as infection control. The numbers below each column indicate the relative protein abundance measured by densitometry and normalized as described in the Materials and Methods.

To further elucidate the temporal effects of the HPIV3 matrix on host protein synthesis, cells were examined at various post-transfection intervals and those expressing HPIV3 matrix trafficking mutants. Host protein synthesis was notably inhibited by the HPIV3 matrix at 24 hpi, with almost 60% inhibition observed at 48 hpi (Figure 1E, lanes 2 and 4). To determine whether the inhibitory effects of the HPIV3 matrix protein were due to its cytoplasmic or nuclear pools, we generated cytoplasmic- and nuclear-resident mutants based on the well-defined nuclear localization signals (NLS) and nuclear export signals (NES) demonstrated for the NiV matrix ^16^. The HPIV3-M mutants exhibited similar distribution patterns to those of NiV-M as assessed by the cytoplasmic-to-nuclear (C/N) ratio of HPIV3-M intensities in transfected HeLa cells (Figure S1). Cytoplasmic-resident mutants (Bp12) from HPIV3 and NiV exhibited a stronger inhibitory effect compared to their wild-type (WT) counterparts (Figure 1F, lanes 3 and 6). Conversely, nuclear-resident mutants (LL) showed a reduced inhibitory impact relative to the WT (Figure 1F, lanes 4 and 7). These findings highlight the importance of matrix localization in modulating host protein synthesis.

Finally, to determine if matrix solely modulates host protein synthesis during HPIV3 infection, we infected cells with either HPIV3 (WT) or its matrix-deleted mutant (HPIV3ΔM) (see methods) and assessed global protein synthesis as before. Unlike WT, HPIV3ΔM failed to inhibit host protein synthesis at 24- and 48 hours post-infection (Figure 1G, lanes 3 and 6). This finding emphasizes the pivotal role of the matrix in regulating host protein synthesis during HPIV3 infection. Taken together, these results clearly demonstrate that the HPVI3 matrix inhibited host protein synthesis during infection.

### Viral matrix enhances viral but suppresses host protein expression at the translational level

Given that the viral matrix can inhibit host protein expression, we aimed to investigate the specific stages of viral and host protein expression targeted by the viral matrix protein, determining whether its effects occur at the level of mRNA transcription or translation. To address this, we engineered a stable cell line expressing a rNiV-NPL replicon (Haas G et al, manuscript in preparation), which encodes essential genes for viral transcription and genome replication in the cell; N, P, L, and a luciferase (Luc) reporter between N and P genes. This replicon allows the production of single-cycle infectious virus-like particles (VLPs) by co-transfecting the NiV matrix, fusion, and receptor-binding proteins (M, F, and RBP). These rNiV-VLPs can infect cells but not produce new virus particles, ensuring that all readouts originate from the initial inoculation. Importantly, the effects of matrix on viral gene and protein expression can be studied by exogenous addition of matrix. Cells expressing increased amounts of NiV-M, followed by infection with rNiV-NPL-VLP, exhibited enhanced Luc activity (Figure 2A, lanes 2-3 compared to Lane 1). Similarly, the HPIV3 matrix, from a different paramyxovirus genus, also enhanced Luc activity (Figure 2A, lanes 4-5 compared to Lane 1). Moreover, increased amounts of matrix protein inhibited host protein synthesis (Figure 2A, lower panel). Analysis of viral RNA expression levels for N, P, and L genes, as well as the viral genome and anti-genome, revealed no significant changes regardless of the presence of matrix proteins. This indicates that the observed enhancement was specific to the translation process and not due to increased mRNA transcription.

**Figure 2.**
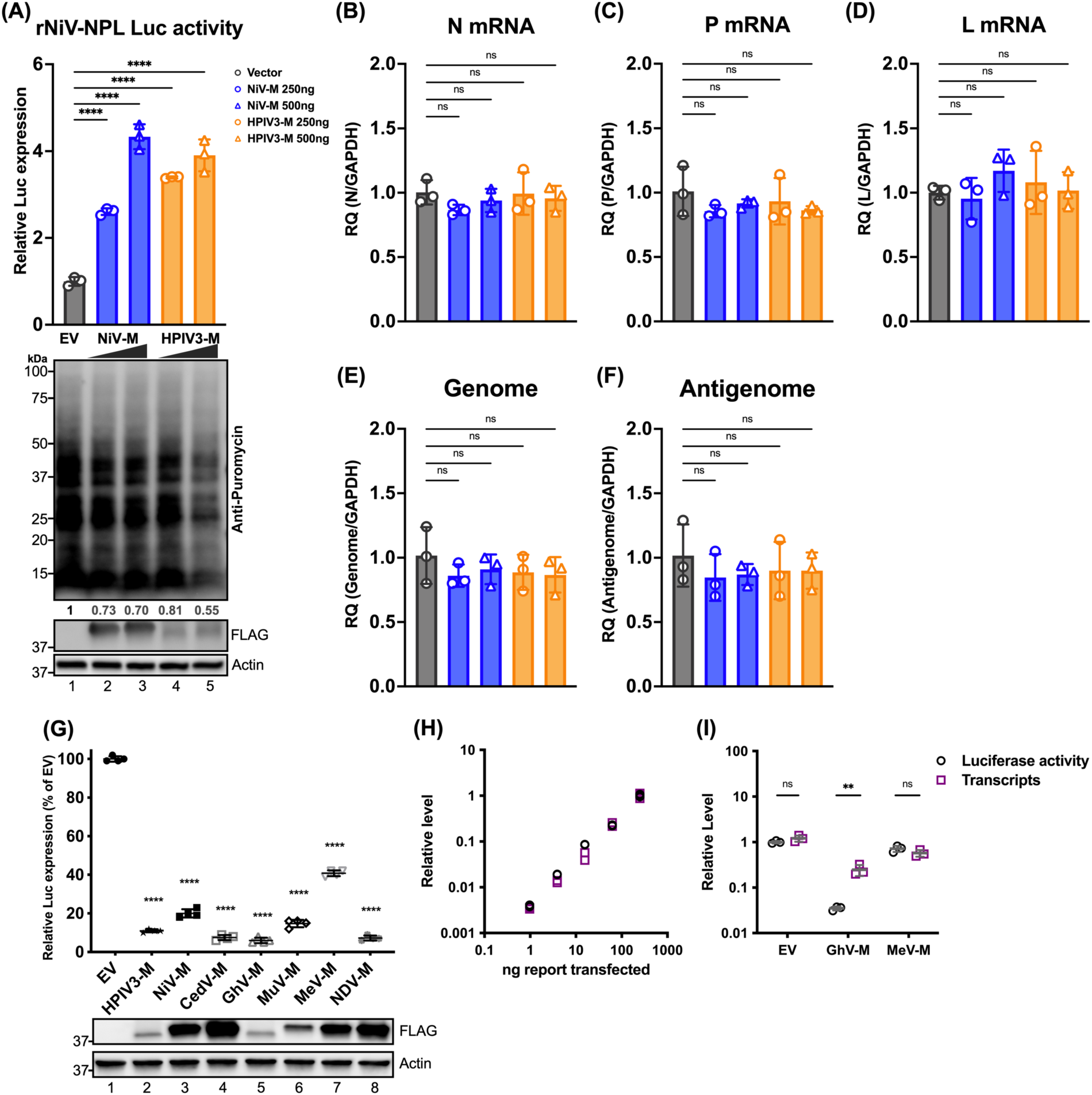
Effects of paramyxoviral-matrix on Nipah-NPL replicon and the expression of cellular splicing-dependent luciferase. (A) HEK-293Ts were transfected with either NiV or HPIV3 matrix for 24 h. Following transfection, cells were inoculated with NiV-NPL replicon virus-like particles (VPL) and incubated for an additional 48 hours. Relative luciferase activity was measured using the Nano-Glo HiBiT system (upper panel). Puromycin-pulsed cells were analyzed by immunoblotting to assess protein expression levels (lower panel). (B-F) Total RNA was extracted from cells treated as in (A) and subjected to RT-qPCR. Relative viral transcript quantity (RQ) was normalized to GAPDH expression. (G) Luciferase activity (RLU) in HEK-293Ts co-transfected with paramyxoviral matrix protein and intron-containing luciferase reporters (Luc-I) for 24 h. RLU was detected using the ONE-Glo system. Expression of FLAG-fused matrixes was analyzed by immunoblotting. Relative luciferase expressions were normalized to EV. (H) A standard curve showing RLUs versus transcripts in HEK-293Ts transfected with varying amounts of Luc-I reporter (250 ng to 0.98 ng, 4-fold serial dilution). RLUs were measured using the ONE-Glo system, and transcript levels relative to 18S rRNA were determined by RT-qPCR. Relative levels were normalized to cells transfected with the maximum amount of Luc-I reporter. (I) Relative levels of RLU and transcripts in HEK-293Ts co-transfected with designated viral matrix and Luc-I reporter. RLUs and transcripts were measured as described in (H). With relative levels normalized to EV. Symbols are data points from biological triplicates. Bar represents the mean ± SD. Statistical significance was determined by one-way ANOVA with Dunnett multiple comparison test. ** *P* <0.01; **** *P* <0.0001; ns, not significant.

To further assess the generalizability PMV matrix inhibiting host protein expression, cells were co-transfected with matrix proteins from multiple PMV genera and an intron-containing Luc reporter (Luc-I) as previously described^46^. The results showed a universal suppression of Luc activity, with the GhV matrix showing a greatest reduction at 90% (Figure 2G, lane 5), while the MeV matrix resulted in a lesser, but still significant, 50% reduction (Figure 2G, lane 7). To determine if this decrease in Luc activity is effectuated at the transcriptional or translational level, we first generated a standard dose-response curve showing a dose-dependent increase in Luc transcripts and (protein) activity levels in cells transfected with increasing amounts of Luc-I (Figure 2H). Using these standard curves, we analyzed the relative levels of Luc transcripts and activity in cells expressing either empty vector (EV), GhV, or MeV matrix, and observed no significant changes between Luc transcript levels and activity in the EV and MeV conditions. However, cells expressing the GhV matrix showed a marked suppression in Luc activity despite only a marginal reduction in Luc transcripts (Figure 2I).

These results collectively suggest that the viral matrix proteins enhance viral protein translation without affecting mRNA transcription and suppress host protein translation at the mRNA translational level.

### HPIV3 matrix promotes ribosome association of viral mRNAs during infection

To assess the impact of HPIV3 infection on host mRNA translation, we performed polysome profiling analysis using HEK-293Ts that were either mock-infected, infected with HPIV3, or transfected with the HPIV3 matrix protein. The polysome profile (Figure S2A) from mock-infected cells (black line) displayed distinct peaks corresponding to the 40S and 60S ribosomal subunits, the 80S monosome, and polysomes, which serve as a baseline. Upon HPIV3 infection (blue line), there was a noticeable accumulation of 80S monosomes, particularly in fractions 6-7, a phenomenon also observed in other viruses such as VSV^47^. This accumulation was further confirmed by densitometry of western blotted proteins across all fractions, which showed increased intensities of small (S6) and large (S7a) ribosomal proteins in their respective cognate fractions (Figure S2B-D). In cells transfected with the HPIV3 matrix protein alone, there was an even more pronounced peak of 80S monosomes, accompanied by a decrease in polysome fractions. Immunoblotting for cEJC components (Figure S2B), including eIF4AIII, Y14, and MAGOH, showed consistent distribution across the ribosomal fractions, aligning with the previous study^48^.

To further characterize the impact of the matrix protein on viral and host translational profiles during infection, we generate a recombinant HPIV3ΔM with the matrix gene substituted by an mCherry reporter, so we could perform polysome transcriptional profiling using isogenic HPIV3 infected cells that differ only in virus expressed M protein. Polysome profile analysis of cells infected with HPIV3 revealed an increase in the 80S monosome pool 48 hours post-infection (Figures 3A, blue line), relative to mock-infected cells (Figure 3A, black line). This confirms the polysome profile presented in Figure S2. However, this increase was abrogated in cells infected with HPIV3ΔM (Figures 3A, red line), indicating a potential role of the matrix protein in monosome accumulation. Next, we isolated free, monosome- and polysome-associated mRNAs and quantified their relative abundance through mRNA sequencing (mRNA-Seq). Sequence alignments to both viral and host genomes revealed that around 10% of the total reads across all fractions were viral, a pattern consistent in cells infected with HPIV3 (Figure 3B), consistent with observations reported for parainfluenza virus 5 (PIV5), another paramyxovirus^49^. However, the amount of viral reads reduced to 4% of total reads in cells infected with the HPIV3ΔM virus, suggesting a decrease in viral mRNA abundance (Figure 3C). Examination of the distribution of viral transcripts across all seven genes showed that HPIV3ΔM did not alter the transcriptional gradient characteristic of paramyxoviruses (Figure 3 D and E). To measure the proportion of viral mRNAs associated with ribosomes, we calculated the ratio of each viral mRNA’s proportion in the ribosome-associated fraction (fractions 7-16) to its proportion in the free fraction (fractions 3-4), which we term “ribosome association efficiency.” Comparative analysis of the ribosome association efficiency between HPIV3 and HPIV3ΔM-infected cells revealed a higher ribosome association ratio for viral transcripts in HPIV3 (Figure 3F and Data S1).

**Figure 3.**
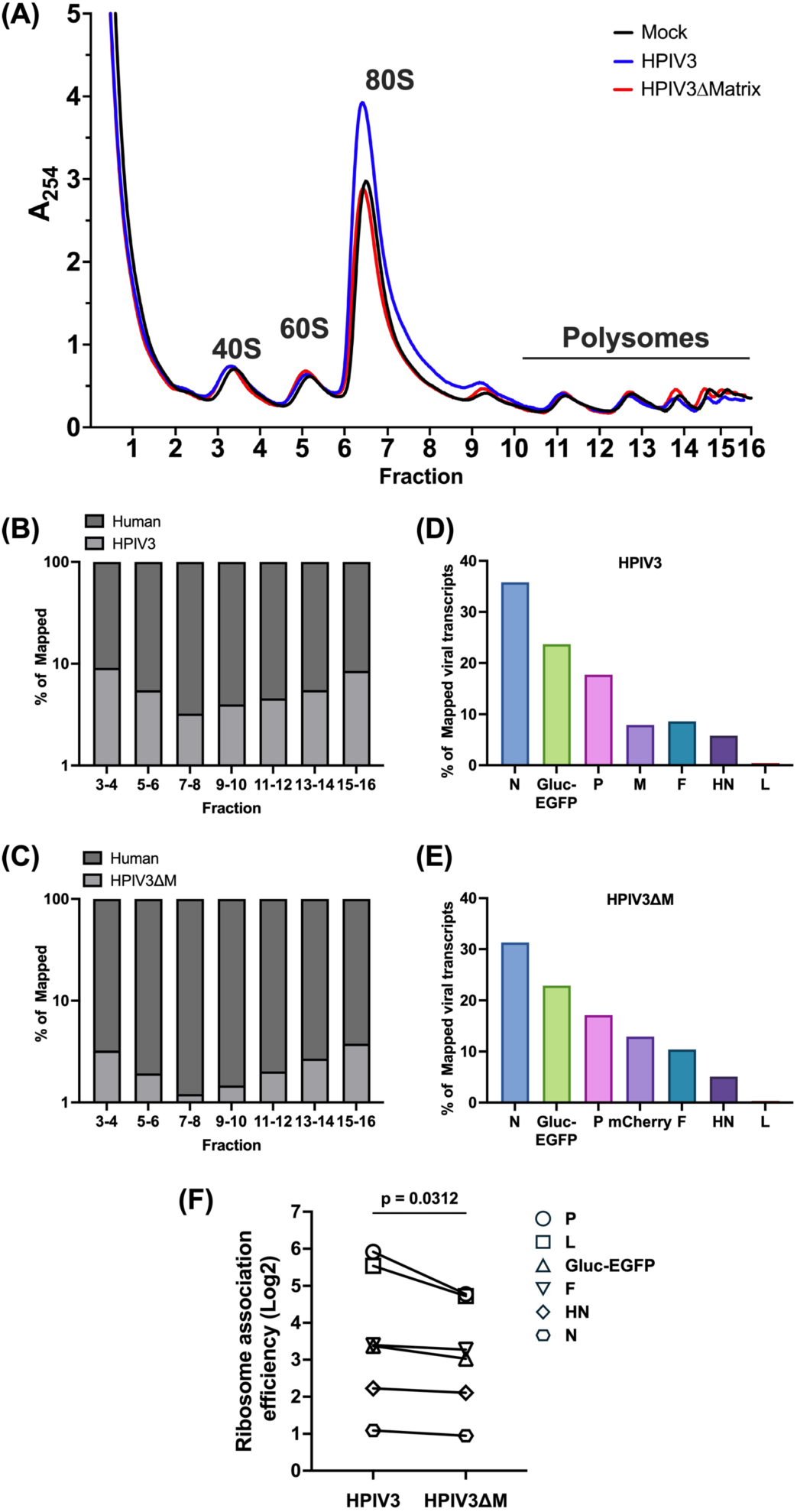
Polysome profile and viral transcript distribution in HPIV3 and HPIV3ΔM infected cells. (A) Polysome profiles of mock-infected (black), HPIV3 infected (blue), or HPIV3ΔM infected (red) HEK-293Ts at 48 hrs post-infection (hpi). HEK-293Ts were infected with HPIV3 or HPIV3ΔM at MOI of 3. Cytoplasmic extracts were prepared at 48 hpi and subjected to sedimentation through a 10-50% sucrose gradient. Absorbance at 254 nm was continuously monitored, and 0.6 ml fractions were collected. Distribution of fragments mapping to (B) human and HPIV3 or (C) human and HPIV3ΔM genome across the sucrose gradient fractions 7 to 16 at 48 hpi. (D-E) Distribution of viral transcript among the seven viral genes of either HPIV3 or HPIV3ΔM infected HEK-293Ts at 48 hpi. The percentage of mapped viral transcripts was quantified using the transcripts per kilobase Million (TPM) metric to normalize for gene length and library size. (F) Comparative analysis of ribosome association efficiency of viral transcripts in HPIV3 and HPIV3ΔM infected cells. Statistical significance was analyzed by Wilcoxon test. * P <0.05.

Taken together, these findings suggest that the matrix potentially enhances viral mRNA translation efficiency by promoting the association of viral transcripts with ribosomes, thereby facilitating more efficient viral translation during infection.

### HPIV3 matrix exhibits a minimal impact on the abundance of individual cellular mRNAs in the monosome and polysome fractions

To determine how the HPIV3 matrix affects the distribution of mRNAs between monosome and polysome fractions, we plotted the transcripts per million (TPM) for each cellular mRNA mapped to the human genome in both fractions. In the monosome fraction, which represents transcripts at the initial stage of translation, the expression abundance of cellular transcripts was similar in both HPIV3 and HPIV3ΔM infections (Figure 4A and Data S2). This similarity indicates that the viral matrix did not significantly alter *relative* cellular gene expression, as shown by the comparable relative transcript ratios between HPIV3 and HPIV3ΔM infections (Figure 4B). In the polysome fraction, representing actively translating transcripts, the expression abundance of cellular transcripts also showed no significant differences between HPIV3 and HPIV3ΔM infections (Figure 4C). The matrix protein altered the abundance of cellular transcripts within a narrow range, with few genes exhibiting more than a 2-fold change in the relative transcript ratio (Figure 4D). Overall, these results suggest that during HPIV3 infection, the presence of the viral matrix protein does not cause significant changes in the translation of cellular transcripts. The distribution of mRNAs between monosome and polysome fractions remains largely consistent, indicating that the matrix protein has a minimal impact on overall cellular gene expression at the translational level.

**Figure 4.**
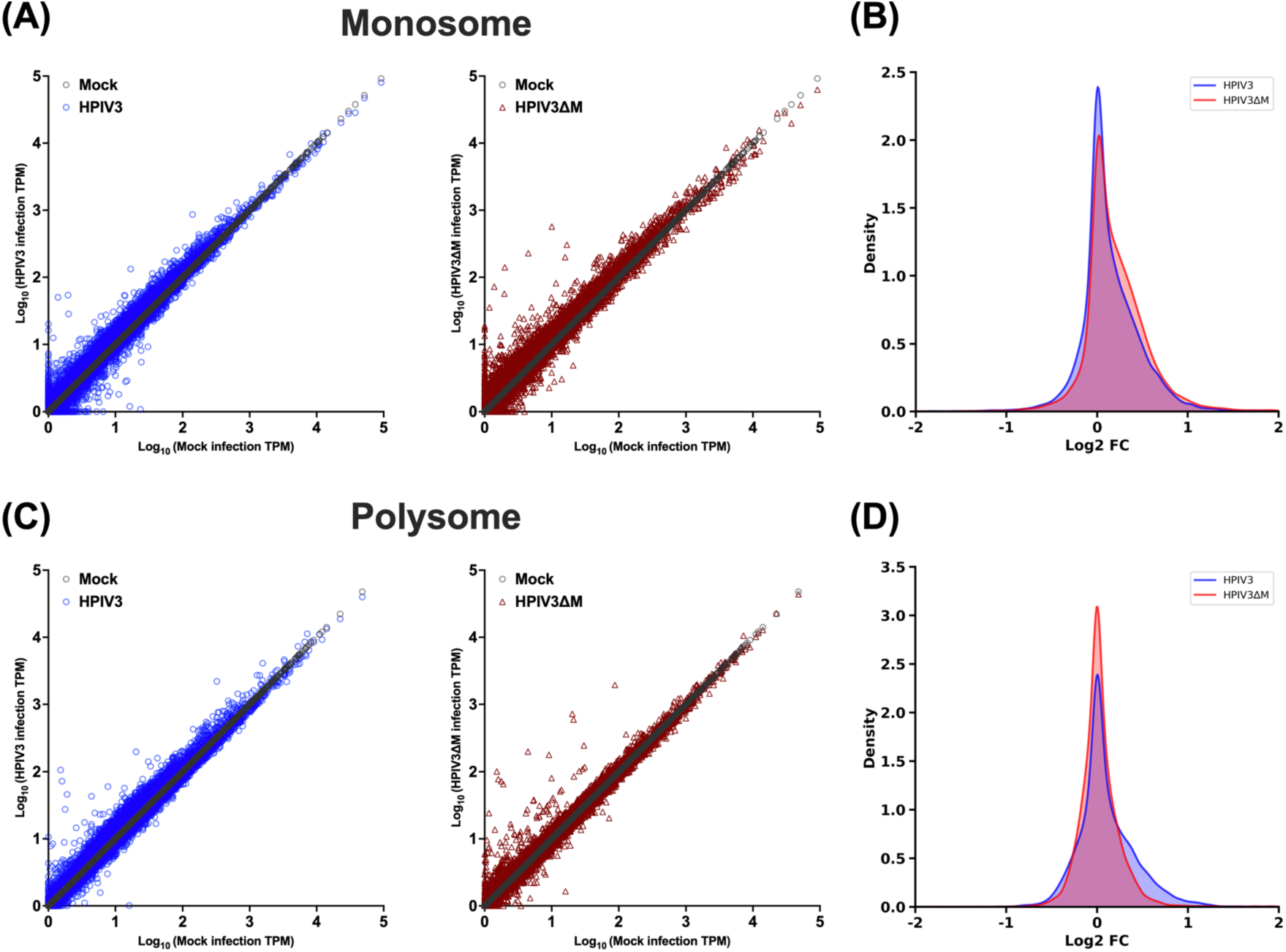
Effects of HPIV3 matrix on the relative abundance of individual cellular mRNAs between monosome and polysome. (A) Scatter plots of transcripts per kilobase Million (TPM) for cellular mRNA transcripts in monosome fraction at 48 hpi. The x-axis graphed unique cellular mRNAs from mock-infected cells, and the y-axis depicted the corresponding TPM values for each mRNA in either mock-infected (gray circles), HPIV3-infected (blue circles) or HPIV3ΔM infected cells (red triangles). (B) Density plots of the log2 fold change in TPM for cellular mRNAs between virus-infected (HPIV3 or HPIV3ΔM) and mock-infected cells in monosome fraction. (C) Scatter plots of TPMs for cellular mRNA transcript in polysome fraction, presented as in A. (D) Density plots of the log 2fold change in TPM in polysome fraction, presented as in B.

### Paramyxoviral-matrix proteins interact with the core components of the exon junction complex

We have found that the matrix disrupts host protein expression at the translation level (Figure 2). To address how the matrix protein disrupts host translation, we sought to identify its potential cellular targets. An inducible 293Ts expressing FLAG-fused NiV-M was generated, enabling the efficient co-purification of NiV-M interacting proteins. The composition of these M-interacting proteins was then characterized using proteomic mass spectrometry, following methods delineated in the previous study^14^. Using the CORUM protein complex database, we discovered a significant enrichment of proteins associated with the exon junction complex (EJC) within the NiV-M interactome (Figure 5A, with a -log10 (adjusted p-value) > 4). The EJC serves as a multifaceted regulator of mRNA biogenesis, including core proteins such as eIF4AIII, Y14, and MAGOH, which were found to interact with NiV-M. To determine whether matrix proteins from various paramyxoviruses interact with the EJC components, we performed co-immunoprecipitation (co-IP) and immunoblot analysis.

**Figure 5.**
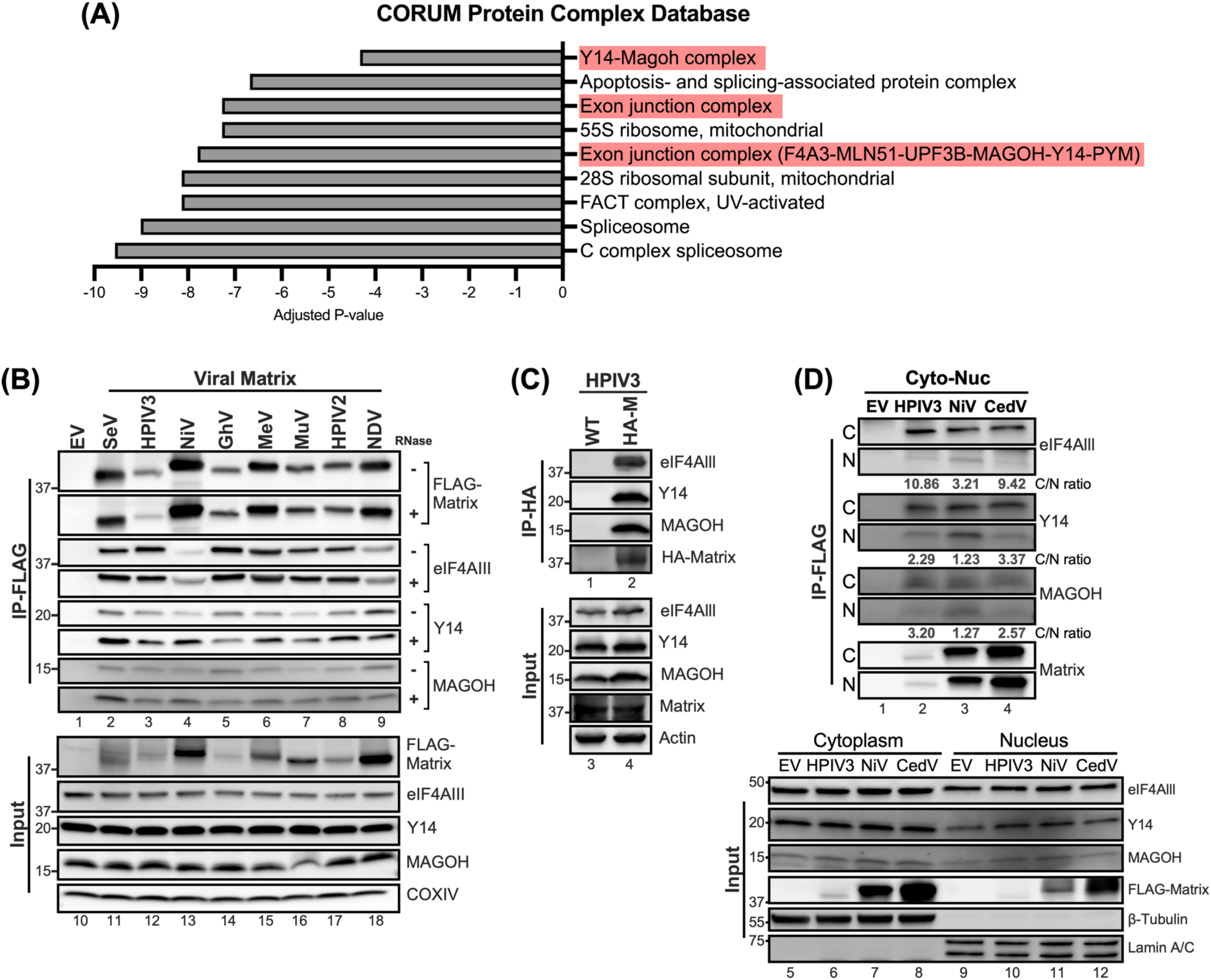
Interactions between Paramyxovirus matrixes and the core components of exon junction complex. (A) Protein complexes enriched in CORUM protein database from matrix interactome identified by MudPIT analysis. Adjusted P-value indicated the significance of the enriched protein complex. (B) HEK-293Ts overexpressing the indicated FLAG-tagged matrix proteins were Immunoprecipitated (+/- RNase) with Anti-FLAG M2 affinity gel after 48 hrs post-transfection. Matrix-bound proteins were analyzed by immunoblotting and endogenous levels of eIF4Alll, Y14, and MAGOH were detected by designated Abs. The amount of input was 5% of total IP lysates. (C) HEK-293Ts were subjected to HA-tag immunoprecipitation following inoculation with HPIV3 containing none or HA-tagged matrix at 0.01 m.o.i at 48 hrs post-infection. (D) Nuclear and cytoplasmic fractions from cells expressing specified FLAG-tagged matrix proteins were subjected to FLAG immunoprecipitation. Subsequent immunoblotting identified interacting proteins. Values below the blots represent the intensity ratios of eIF4AIII, Y14, and MAGOH from cytoplasm to nucleus. β-Tubulin and Lamin A/C served as cytoplasmic and nuclear fraction markers, respectively. IP, immunoprecipitation. IB, immunoblot.

Cells expressing FLAG-fused matrix from 8 different paramyxoviruses were subjected to immunoprecipitation using Anti-FLAG M2 affinity beads. The results revealed that the core components of EJC (cEJC), the eIF4AIII, Y14, and MAGOH were all present in matrix precipitants (Figure 5B, lane 2-9), but not in the FLAG precipitant (vector control, lane 1). To ensure that the interaction between matrix and cEJC is RNA-independent, we conducted a similar experiment with the addition of RNase A to each reaction. The amount of cEJC in matrix immunoprecipitants was unchanged, or in some cases, even increased (Figure 5B, RNase A +), suggesting the RNA-independent association between the paramyxoviral matrix proteins and the cEJC components.

To ensure that matrix-cEJC interactions were not an artifact of matrix overexpression during transient transfections, we sought to validate the interaction between the matrix and the cEJC during infection. We infected 293T cells with either wild-type HPIV3 (WT) or a previously characterized isogenic counterpart expressing an HA-tagged matrix protein (HPIV3_HA-M)^50^ and conducted HA-IP at 48 hpi. Notably, cEJC components were detected exclusively in HA-coprecipitates from cells infected with the HPIV3_HA-M, but not in those from WT-infected cells (Figure 5C, lanes 1 and 2), thereby validating the specific interaction between the matrix and cEJC in the context of HPIV3 infection. Having verified the interaction between matrix and cEJC, we next wanted to assess where this matrix-cEJC interaction occurred as both matrix and cEJC components are known to undergo nuclear-cytoplasmic trafficking. 293Ts expressing FLAG-tagged matrix from HPIV3, NiV, and CedV were subjected to cytoplasmic-nuclear fractionation. FLAG-IP was then employed to determine the interaction dynamics of the matrix proteins and cEJC within these fractions. Analysis of the input (Figure 5D, lanes 5 to 12) revealed that the cEJC components were more abundantly present in the cytoplasm compared to the nucleus. Additionally, FLAG-tagged matrix proteins from HPIV3, NiV, and CedV showed a relatively even distribution between the cytoplasmic and nuclear fractions. Following the IP, results revealed that the band intensities for cEJC co-IPed with the matrix were markedly stronger in the cytoplasmic fractions compared to the nuclear (Figure 5D, lanes 2 to 4). Quantification of these bands yielded cytoplasm-to-nucleus intensity ratios consistently greater than 1 (Figure 5D, C/N ratio), suggesting a preferential interaction within the cytoplasm. These results collectively indicate that the matrix-cEJC interaction predominantly occurs within the cytoplasm, highlighting a potential mechanism by which the matrix protein disrupts host translation.

### HPIV3 infection perturbs the subcellular distributions of core EJC components

To potentiate its own replication, viruses have evolved strategies to co-opt or antagonize functions of cellular proteins, including altering their expression levels and localization. Although we did not detect a significant change in the abundance of cEJC during HPIV3 infection (Figure 6A), we suspected that HPIV3 infection might alter the localization of these proteins as described in previous studies with flaviviruses^51–53^. We performed cytoplasmic-nuclear fractionation in HPIV3-infected cells and monitored the relative abundance of cEJC in the cytoplasmic and nuclear fractions (Figure 6B). We observed an increased abundance of eIF4AIII, Y14, and MAOGH in the cytoplasmic fraction, with a concomitant decrease in the nucleus fraction during infection, while the total amount of cEJC remained unchanged in the infected whole cell lysates. Densitometric quantification of the cognate band intensities showed that the cytoplasmic: nuclear ratios of eIF4AIII, Y14, and MAOGH increased 3-5 fold at 24 hpi. To gain further insights, we visualized the localization of the HPIV3 matrix and cEJC components in HPIV3-infected cells via confocal microscopy. We first address whether the HPIV3 matrix exhibits a similar trafficking behavior to the NiV matrix, as previously reported^14,16^. The cytoplasmic matrix translocated into the nucleus at 12-16 hpi and then returned to the cytoplasm by 24 hpi (Figure S3A). This was further confirmed by the increased cytoplasmic-to-nuclear (C/N) ratio of HPIV3-M intensities (Figure S3B). At 24 hpi, the HPIV3 matrix was observed at the plasma membrane of infected cells, consistent with its role in the budding process of mature virions (Figure 6C-E). Compared to the mock, HPIV3 infection appeared to disrupt the distribution of cEJC components, leading to their accumulation in the cytoplasm, consistent with our previous fractionation results. More importantly, we observed partial, rather than complete, colocalization of matrix and cEJC components in cytoplasmic puncta (Figure 6C-E, orthogonal projections), suggesting that viral matrix may modulate the cytoplasmic function of the EJC, such as mRNA translation. To further confirm that the HPIV3 matrix is responsible for the accumulation of cEJC in the cytoplasm, we monitored the distribution of the eIF4AIII in cells infected with HPIV3ΔM. Compared to the HPIV3, the ΔM virus lost the ability to redistribute the eIF4AIII (Figure 6F), as indicated by the low C/N ratio of eIF4AIII intensities.

**Figure 6.**
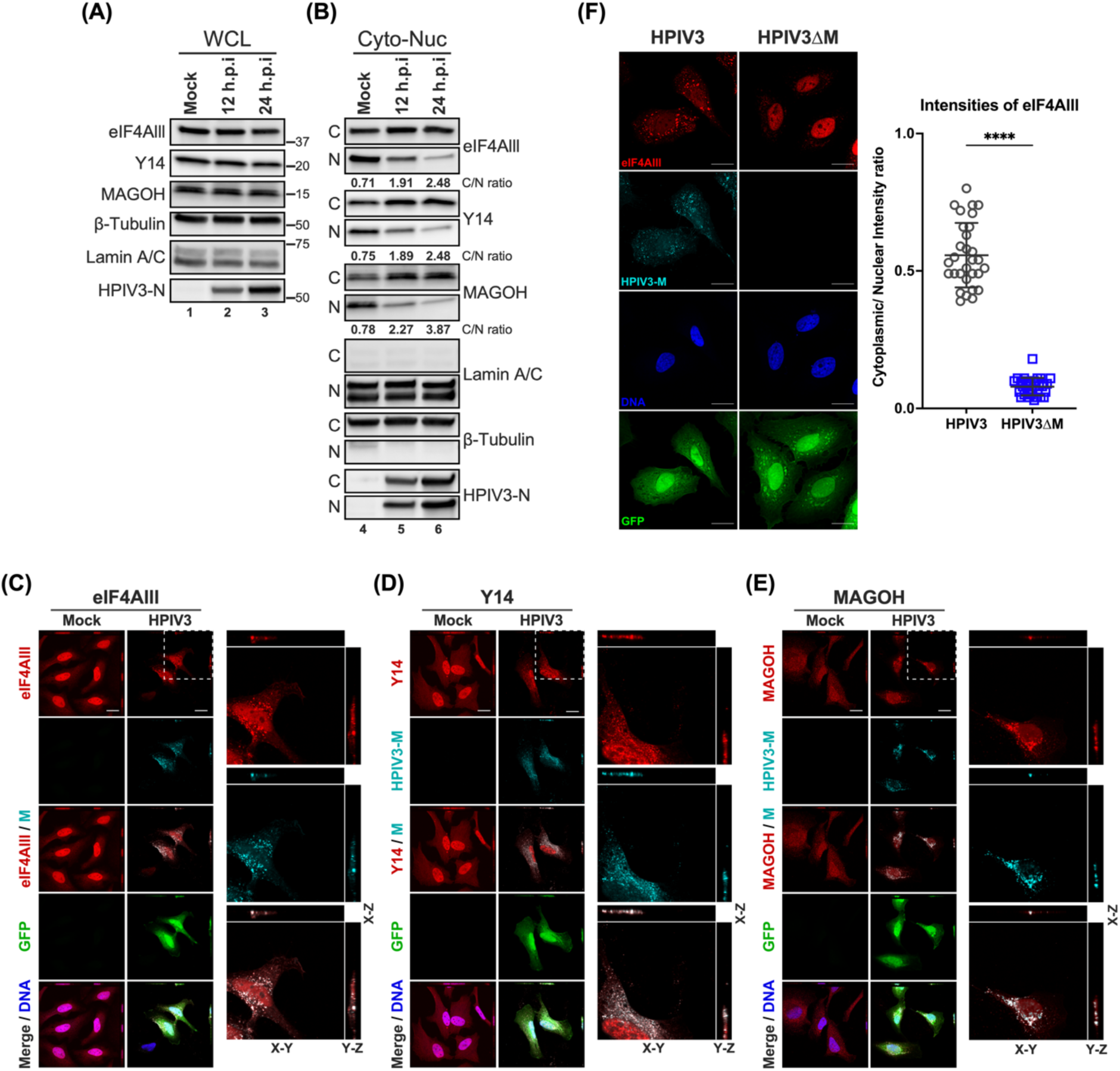
Subcellular localization of core EJC in HPIV3 infected HeLa cells. (A-B) Immunoblotting analysis of whole cell lysates, cytoplasmic, and nuclear fractions from HPIV3-infected HeLa cells at 12- and 24-hours post-infection (hpi). The levels of eIF4AIII, Y14, and MAGOH were examined, with β-tubulin and Lamin A/C serving as markers for the purity of cytoplasmic and nuclear fractions, respectively. The ratios below the blots indicate the relative intensities of eIF4AIII, Y14, and MAGOH from the cytoplasm to the nucleus. (C-E) XYZ planes of 3D confocal micrographs depicted HeLa cells at 24 hours post-infection with HPIV3 at m.o.i of 5. Cells were fixed and stained with (C) anti-eIF4AIII, (D) anti-Y14, or (E) anti-MAGOH antibodies (red), and anti-HPIV3-M antibody (cyan) to label the viral matrix protein. Nuclei were counterstained with Hoechst (blue), and GFP fluorescence indicates HPIV3 infection. Enlarged orthogonal projections of the infected cells (white dashed line) are shown on the right, displaying the EJC protein, HPIV3-M, and the merged channels. Scale bars represent 20 µm. (F) Left: HeLa cells infected with either HPIV3 or HPIV3ΔM at an m.o.i. of 5 were fixed at 24 hours post-infection and stained with anti-eIF4AIII (red), anti-HPIV3-M antibodies (cyan), Hoechst for nuclei (blue), and GFP fluorescence indicates HPIV3 infection. Representative fields of cells for each condition are shown. Right panel: Quantification of cytoplasmic/nuclear eIF4Alll intensity (C: N) ratios was performed on 30 individual cells, as described in Materials and Methods. Statistical significance was analyzed by unpaired t-test. **** P <0.0001.

Taken together, these findings demonstrate that the HPIV3 matrix induces the redistribution of cEJC components to the cytoplasm, leading to their partial cytoplasmic colocalization with the matrix. cEJC typically docks on nascent mRNAs after splicing, serving as a platform for factors involved in mRNA export and enhancing mRNA translation efficiency. These components are usually recycled back into the nucleus to continue their role in mRNA biogenesis; this co-localization in the cytoplasm might restrict the cellular functions of the cEJC, potentially impacting host mRNA translation during infection.

### Enhanced viral gene translation through eIF4AIII knockdown and matrix expression

To ascertain the role of cEJC components, specifically eIF4AIII, in the stages of viral replication, we utilized the rNiV-NPL replicon system described in Figure 2. The knockdown of eIF4AIII was employed to simulate the restriction of cellular functions by the viral matrix, allowing us to closely examine the impact of eIF4AIII on viral mRNA transcription and translation. Cells were co-transfected with either negative control siRNA (siNC) or siRNA pool targeting eIF4AIII (sieIF4AIII) along with an empty vector (EV) or plasmids expressing increased amounts of NiV-M or HPIV3-M for 24 hrs. The cells were then infected with rNiV-NPL-VLP, and Luc activity was measured to indicate the level of viral gene translation. Compared to siNC, the knockdown of eIF4AIII enhanced Luc activity in the rNiV-NPL-VLP infected cells (Figure 7A upper panel, lane 2 vs. lane 1); knockdown of eIF4AIII also slightly inhibited host protein synthesis as indicated by the western blot (lower panel, Lane 2 vs. Lane 1). In the eIF4AIII knockdown background, the addition of NiV or HPIV3 matrix further enhanced Luc activity in a dose-dependent manner (Lanes 3-6 vs. Lanes 1 and 2). The RNA levels of the viral genes and genomes showed minor changes (less than 2-fold) in cells expressing EV or matrix proteins in the eIF4AIII knockdown background compared to the cell expressing EV in the NC knockdown background (Figure 7B-7F), which was not as pronounced as the changes observed in Luc activity. These findings suggest that the partial knockdown of eIF4AIII leads to a slight reduction in host protein synthesis but enhances viral gene translation, suggesting a redistribution of translational resources to viral mRNAs. Furthermore, the expression of the viral matrix in cells with partial eIF4AIII knockdown further restricts the function of the remaining eIF4AIII, enhancing viral gene translation. Thus, we propose that the viral matrix enhances viral gene translation by restricting the cellular function of eIF4AIII.

**Figure 7.**
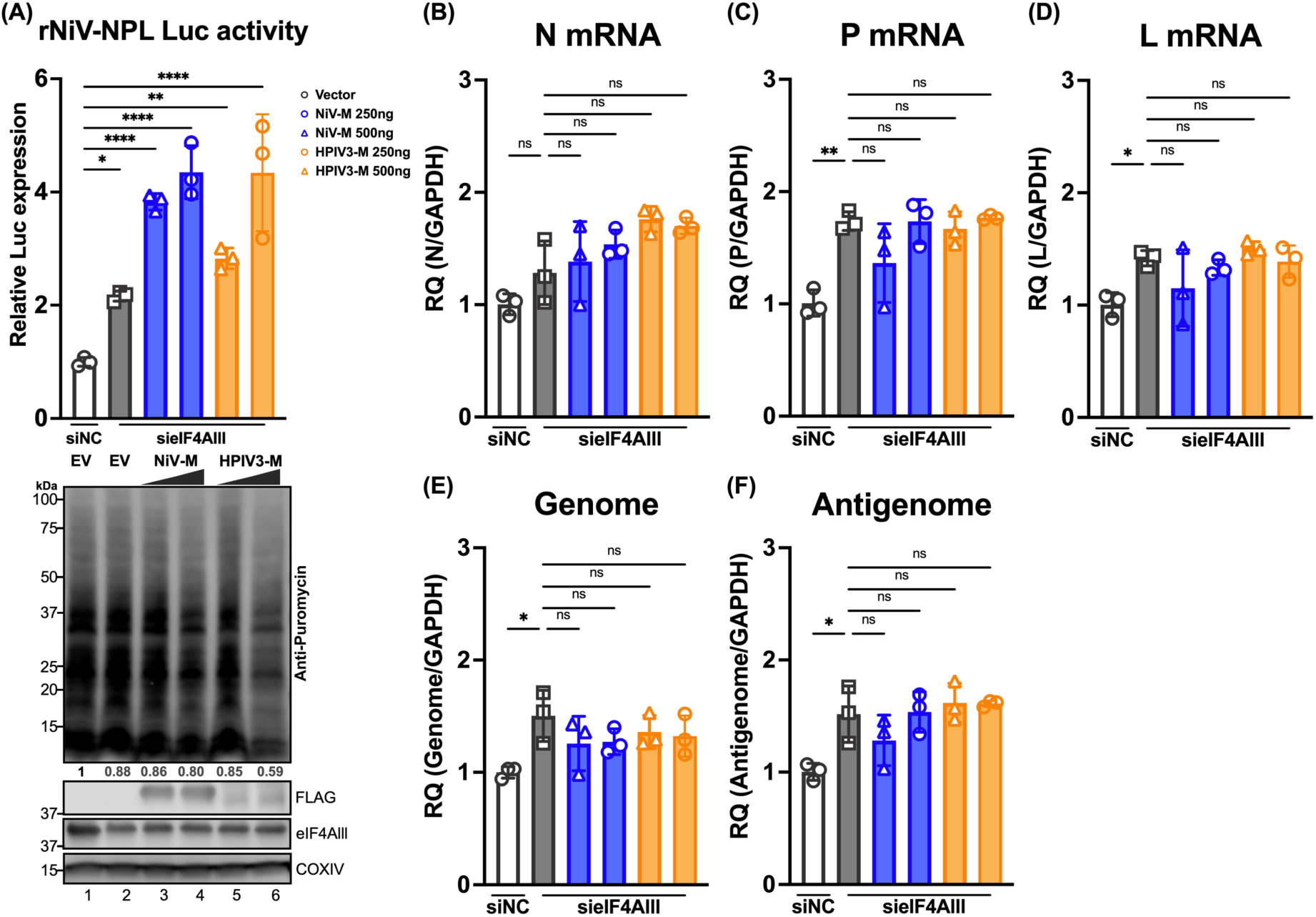
Effects of paramyxoviral-matrix on Nipah-NPL replicon in eIF4Alll knockdown cells. (A) HEK-293T cells were co-transfected with either control siRNA (siNC) or siRNA pool targeting eIF4AIII (sieIF4AIII) along with an empty vector (EV) or plasmids NiV-M or HPIV3-M 24 h. Following transfection, cells were inoculated with rNiV-NPL replicon virus-like particles (VPL) and incubated for an additional 48 hours. Relative luciferase activity was measured using the Nano-Glo HiBiT system (upper panel). Puromycin-pulsed cells were analyzed by immunoblotting to assess protein expression levels (lower panel). (B-F) Total RNA was extracted from cells treated as in (A) and subjected to RT-qPCR. Relative viral transcript quantity (RQ) was normalized to GAPDH expression. Symbols are data points from biological triplicates. Bar represents the mean ± SD. Statistical significance was determined by one-way ANOVA with Dunnett multiple comparison test. * *P* <0.05; ** *P* <0.01; *** *P* <0.001; **** *P* <0.0001; ns, not significant.

### Core EJC depletion can enhance paramyxovirus replication

Building on our findings that partial knockdown of eIF4AIII enhances viral gene translation when viral matrix proteins are expressed, we further explored the functional role of core EJC components in authentic paramyxovirus replication. To do this, we utilized siRNA-mediated knockdown (KD) targeting cEJC components: eIF4AIII, Y14, MAGOH, as well as a non-targeting siRNA control. These siRNAs were transfected in 293Ts for 48 hrs prior to infection with a panel of GFP-reporter paramyxoviruses encompassing the 3 major subfamilies of paramyxoviruses, including Human Parainfluenza Virus 3 (HPIV3) and Cedar virus (CedV) (*Orthoparamyxovirinae*), Mumps virus (MuV) (*Rubulavirinae*), or Newcastle Disease Virus (NDV) (*Avulavirinae*). The KD of cEJC components led to an increase by up to 8-fold in the number of infected (GFP-positive) cells across all tested paramyxoviruses (Figure S4). The viral titers were also assessed at 24, 48, or 72 hours of post-infection and demonstrated a more marked increase in cells with cEJC KD, with varying magnitudes of enhancement. Notably, MAGOH KD in HPIV3-infected cells led to a two-log increase in viral titer 48 hpi compared to the non-targeting control (Figure 8A); a comparable trend was also observed in CedV-infected cells with MAGOH depletion (Figure 8B). Conversely, cells with Y14 depletion showed the most pronounced titer increase for MuV and NDV at 72 hours post-infection (Figure 8C-D), suggesting that the observed effects might be influenced by the unique replication dynamics inherent to each virus. To determine if the effect of cEJC KD was specific to paramyxoviruses, we further examined the effects of cEJC on the replication of other RNA viruses such as influenza A, enterovirus D68, and SARS-CoV-2. cEJC depletion did not universally augment the replication of these viruses (Figure 8E-G). In the case of influenza A virus replication, eIF4AIII KD slightly decreased viral titers, consistent with previous reports^54^ implicating eIF4AIII as a positive regulator of IAV replication.

**Figure 8.**
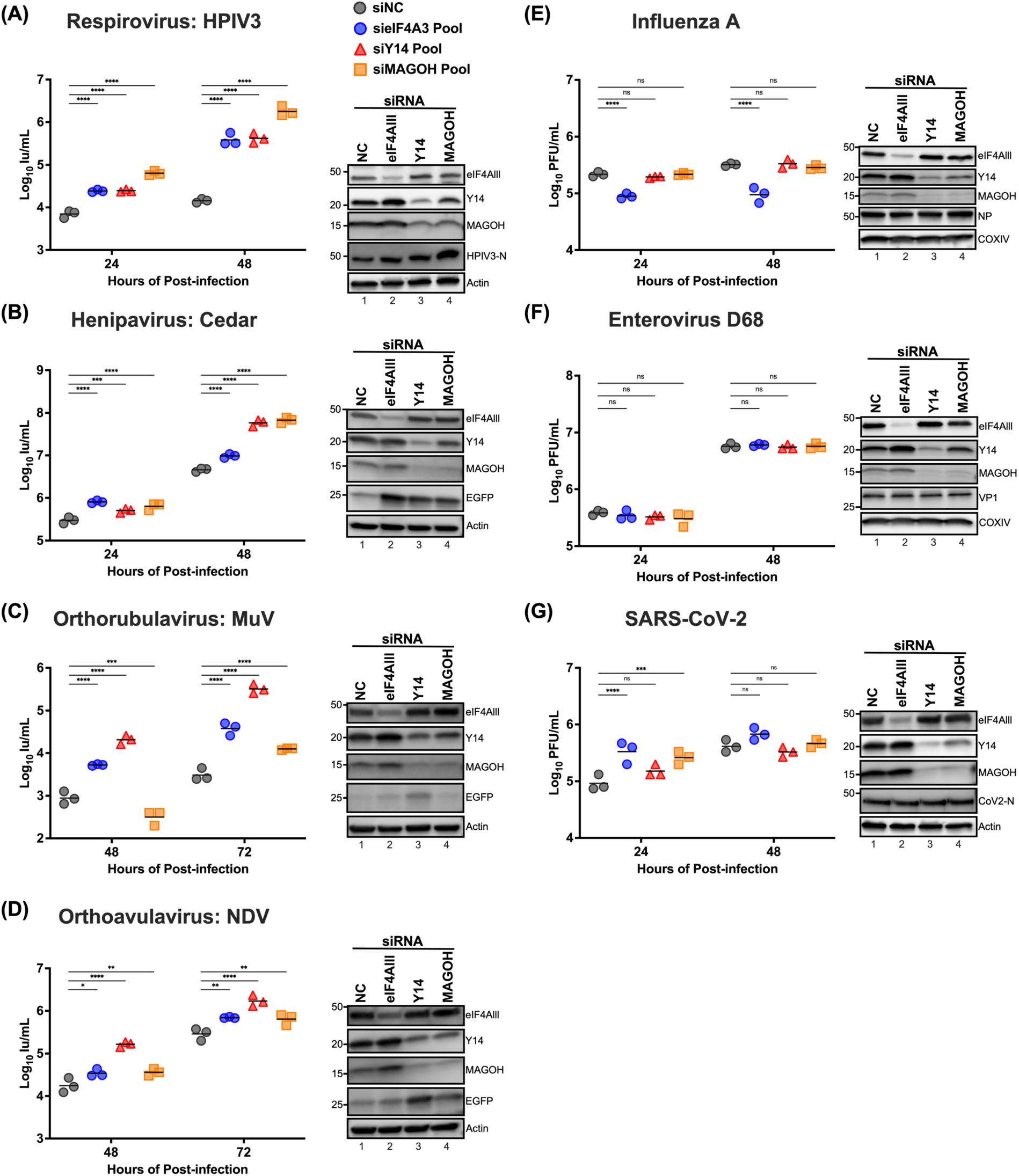
Effects of the exon junction complex on Paramyxovirus, Influenza A, Enterovirus D68, and SARS-CoV2 replication. HEK-293T cells were transfected with siRNA pools targeting eIF4A3, Y14, MAGOH, or non-targeting control siRNAs (NC), respectively. At 48 hrs post-transfection, cells were inoculated with the designated virus (A) HPIV3, (B) Cedar, (C) MuV, (D) NDV, (E) Influenza A (A/WSN/1933), (F) Enterovirus D68, and (G) SARA-CoV2 at a multiplicity of infection (m.o.i.) of 0.01. The titers of infectious supernatants were determined on Vero-CCL81 cells using a 10-fold serial dilution at the indicated time points. For each virus, the expression levels of endogenous eIF4AIII, Y14, and MAGOH, along with infection control for viral protein or EGFP reporter, were analyzed by immunoblotting; results shown beside each panel confirm the knockdown of target proteins and validate virus infection. Symbols represent the data points from biological triplicates. Bars represent the mean of the triplicates. Statistical significance was determined by two-way ANOVA with Dunnett multiple comparison test. * *P* <0.05; ** *P* <0.01; *** *P* <0.001; **** *P* <0.0001; ns, not significant. Immunoblottings are shown beside each to determine the knockdown of target proteins and controls for virus infection.

Collectively, our results indicate that core EJC depletion can significantly and specifically enhance the replication of paramyxoviruses, with the extent of enhancement varying among different viruses.

## DISCUSSION

In this study, we uncovered a novel mechanism by which the HPIV3 matrix manipulates host cell machinery to promote viral translation. Our findings demonstrate that the HPIV3 matrix significantly inhibits host protein synthesis while enhancing viral protein translation, primarily through increased ribosome association with viral mRNAs. Remarkably, we identified a previously unrecognized interaction between the paramyxoviral matrix and core components of the exon junction complex (EJC), predominantly occurring in the cytoplasm. This interaction leads to altered subcellular distribution of EJC components during infection. Functional studies revealed that depletion of EJC components enhances viral gene translation and replication specifically for paramyxoviruses. These results illuminate a strategy employed by paramyxoviruses to subvert host cellular processes, offering new perspectives on virus-host interactions.

The paramyxoviral matrix is known as a structural component localized in the cytoplasm. Several studies, however, have uncovered the nuclear sojourn of the matrix during MeV and NDV infections and transfections, suggesting a role beyond structural functions. One of these nuclear functions is the suppression of host mRNA transcription, which significantly impacts the host’s antiviral response. Despite these findings, the effect of the paramyxoviral matrix on host translation remains unexplored. Using puromycylation to monitor newly synthesized proteins, we have discovered that paramyxovirus infection results in a partial shutdown of host protein synthesis, a phenomenon not previously reported (Figure 1B-C). Our findings indicate that this inhibitory effect on host protein translation is solely attributed to the viral matrix (Figure 1D and 1G). Further investigation into the stages of protein synthesis targeted by the matrix revealed that the disruption occurs at the mRNA translation level (Figure 2). This discovery underscores the significance of the matrix’s localization in hindering host translation. The matrix protein’s ability to inhibit host protein synthesis at this specific stage highlights its strategic role in manipulating the host’s translational machinery to favor viral replication.

To characterize the cellular localization of the HPIV3 matrix, sequence alignment is performed in HPIV-M with NiV-M whose NLS and NES sequences are well defined in the previous studies. Putative NLS (Mbp1/2) and NES (L106A L107A) mutants were introduced into GFP-fused HPIV3-M to visualize their localization in expressing cells. Indeed, Cells expressing WT or mutants all resemble similar distributions as we previously observed in the NiV-M (Figure S1)^14,16^. Furthermore, the localization of HPIV3 during infection (Figure S3) reveals that HPIV3-M has the same trafficking behavior as previously demonstrated in NiV-infected HeLa cells^16^. Similar to the NiV-M, HPIV3-M translocases into the nuclear at early time points (12 to 16 hrs) and distributes into the cytoplasm and the plasma membrane at later time points (20 to 24 hrs). The distribution of HPIV3-M in the cytoplasm might explain several observations in this study:

1. The subcellular localization is critical for the matrix to modulate host translation since the NES mutant of both HPIV-M and NiV-M shows reduced levels of the inhibitory effect by matrix (Figure 1F).
2. Paramyxoviral matrixes suppress host protein synthesis at least, at the mRNA translation level – since mRNA translation also occurs in the cytoplasm (Figure 2A, G, and H).
3. The matrix-cEJC interaction and co-localization are mainly detected in the cytoplasm (Figure 5D and 6C-6E). These observations emphasize the importance of the cytoplasmic-resident matrix for modulating host translation. The expression of wild-type or mutant HPIV3-M alone also shows the cytoplasmic retention of endogenous eIF4AIII at similar levels (Figure S5). However, the pattern of eIF4AIII cytoplasmic retention appears slightly different compared to HPIV3 infection (Figure 6F), as more cytoplasmic puncta are observed during infection. This difference may be due to the matrix co-opting with other viral factors to form liquid-liquid phase separation puncta.

EJC serves as a multifaceted modulator of mRNA biogenesis, it travels with mRNA across different cellular landscapes from pre-mRNA splicing to downstream, posttranscriptional processes such as mRNA export, mRNA localization, translation, and nonsense-mediated mRNA decay (NMD)^55–57^. While NMD is known to impact the infection dynamics of flaviviruses^51–53^, and viruses have evolved various mechanisms to evade or hijack NMD^58,59^, it remains unclear whether HPIV3 infection exerts any interference on NMD activity. To interrogate the effect of HPIV3 infection on NMD activity, we assessed the levels of endogenous NMD targets as delineated in a previous study^53^, which includes SC35, GABARAPL1, ASNS, and CARS. We found that the expression of these NMD targets increases at 48 hours post-infection, suggesting that NMD is impeded by HPIV3 infection (Figure S6). To further decipher whether this suppression is a consequence of the matrix-cEJC interaction, we transfected cells with vectors expressing either GFP or HPIV3-M to monitor the effects of the matrix on NMD activity. To our surprise, the overexpression of HPIV3-M did not disrupt NMD activity, suggesting that the observed inhibitory effects on NMD may be attributed to other viral factors, not the matrix (Figure S7).

The differential impact of cEJC depletion on the replication of paramyxoviruses versus Influenza, SARS-CoV-2, and EVD68 offers intriguing insights into the unique strategies employed by these viruses to hijack host translation machinery. Understanding these mechanisms explains why cEJC depletion benefits paramyxovirus replication but not the other viruses. SARS-CoV-2 NSP1 inserts into the mRNA entry channel on the 40S ribosomal subunit, blocking host mRNA loading for translation initiation. It also interacts with the mRNA export receptor NXF1-NXT1 and NPC proteins (Nup358, Nup214, Nup153, Nup62), preventing proper mRNA export and fostering viral mRNA translation in the presence of the SL1 5’UTR hairpin^29,60,61^. Influenza virus takes over host translation by cap-snatching cellular pre-mRNAs, inhibiting pre-mRNA polyadenylation, and preferentially translating viral mRNAs through sequences in the viral mRNA 5’-UTR^33–36,62,63^. It also retains cellular mRNAs in the nucleus by inhibiting pre-mRNA splicing and blocking mRNA nucleocytoplasmic transport^64–66^. Enterovirus hijacks host translation by using viral proteinase 2A to cleave eIF4E, leading to a global shutdown of Cap-dependent translation, and redirecting ribosomes to viral mRNAs via IRES elements^38,39^. The aggressive global shutoff mechanisms of SARS-CoV-2, Influenza, and EVD68 might overshadow any potential benefits from cEJC depletion, rendering any changes unobservable. In contrast, the paramyxoviral matrix might not completely antagonize cEJC’s cellular function, as evidenced by partial host translation suppression and colocalization of matrix and cEJC (Figure 1 and Figure 6C-E). During paramyxovirus infection, both cellular and viral mRNA are capped and polyadenylated, meaning the virus needs to outcompete for translation resources. The key difference is that host mRNA is docked with EJCs at exon-exon junctions during the pioneer round of translation, while viral RNA is not. Consequently, the depletion of cEJC might inhibit the translation of EJC-bound cellular mRNA, thereby redirecting translational resources to viral mRNA. Moreover, the depletion of cEJC mimics the matrix interaction within the cytoplasm, sequestering EJC-bound cellular mRNA away from the translation apparatus. Additionally, the paramyxoviral matrix might selectively enhance viral mRNA translation, further facilitating the virus’s ability to hijack the host’s translation machinery. Future work uncovering the means for matrix specificity on viral mRNA may unearth novel strategies for selective translational control during viral infection.

## ACKNOWLEDGMENTS

B.L. acknowledges the Ward-Coleman estate for endowing the Ward-Coleman Chairs at the Icahn School of Medicine at Mount Sinai. This work was supported by NIH Grant R01 AI125536 to B.L. C.T.H. and H.P.C acknowledges the postdoctoral fellowship from the National Science and Technology Council, Taiwan. C.S. acknowledges funding from NIH T32 AI07647. G.D.H. was supported by a National Science Foundation Graduate Research Fellowship (NSF GRFP1842169). P.T. was supported by a Canadian Institutes of Health Research fellowship (CIHR 359156). R.W. was supported by an EMBO long-term postdoctoral fellowship (ALTF 628-2015). We thank the members of the Lee lab for their experimental assistance and feedback on the project.

## AUTHORS CONTRIBUTIONS

C-T.H. P.T. and B.L. designed research

C-T.H., G.D.H., R.E.W., H.P.C, S.K., A.P and J.A.W. performed research

G.D.H. contributed new reagents

C-T.H., C.S.S., and B.L. analyzed data C-T.H. and B.L. wrote the paper.

## STAR METHODS

### KEY RESOURCE TABLE

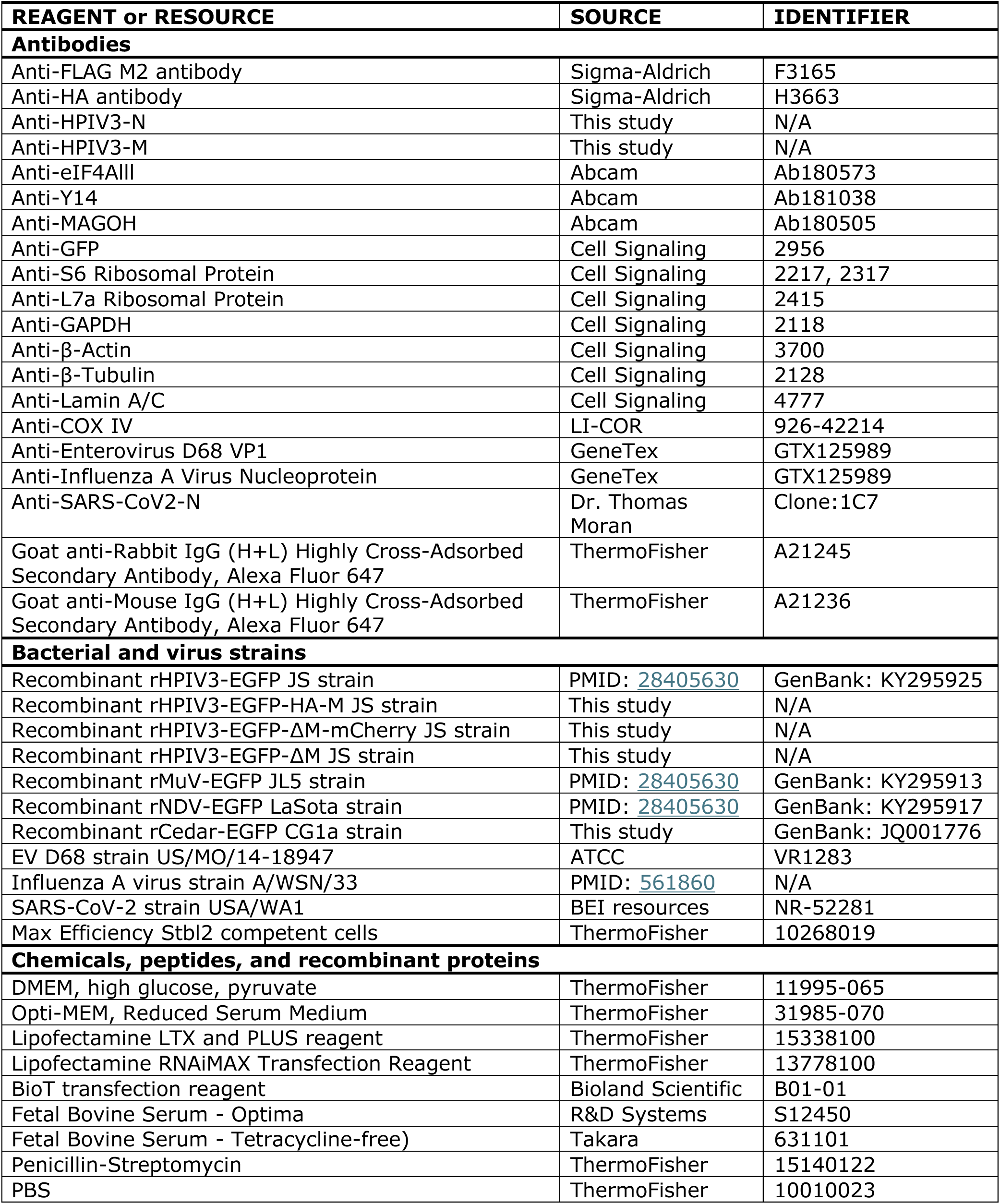
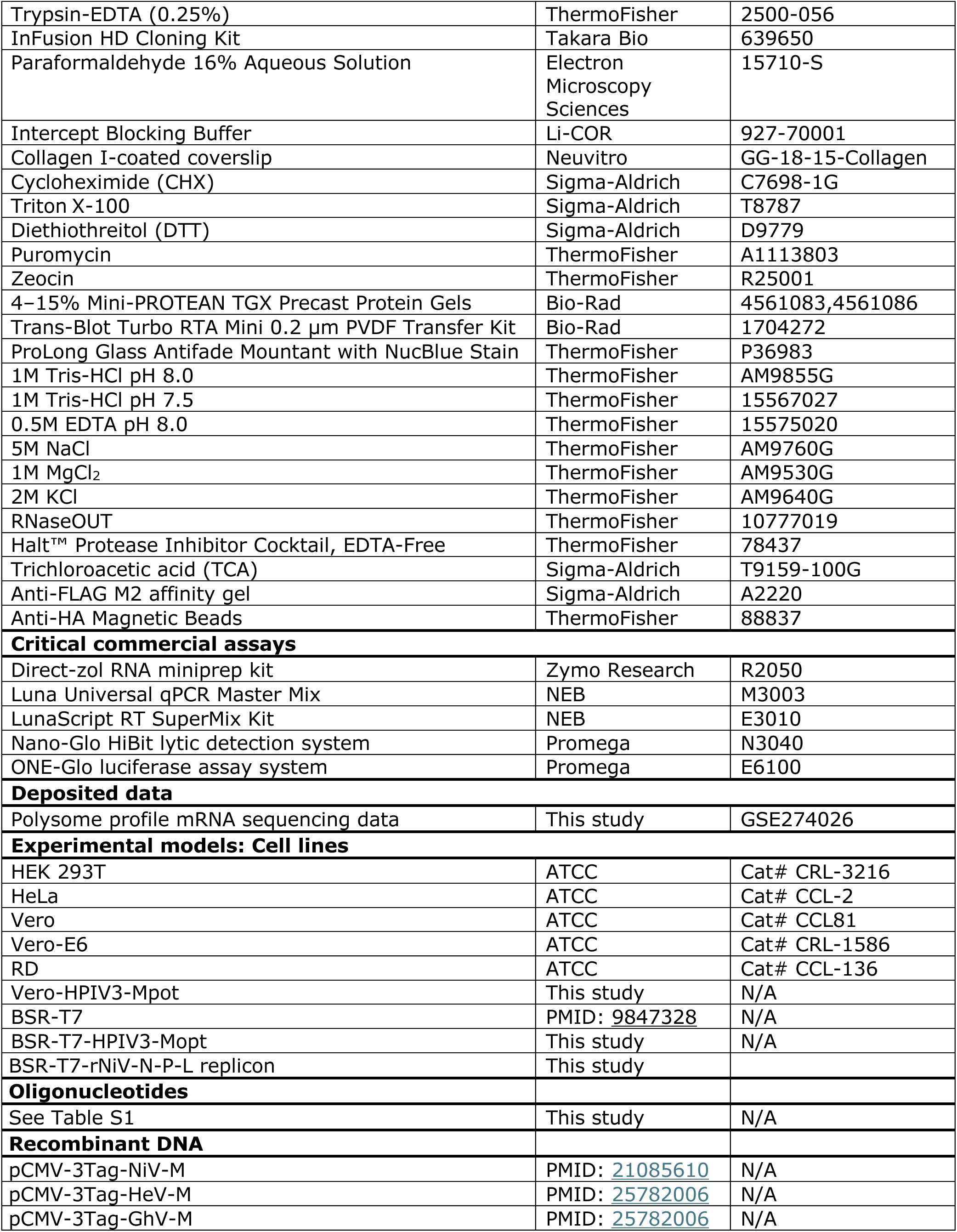
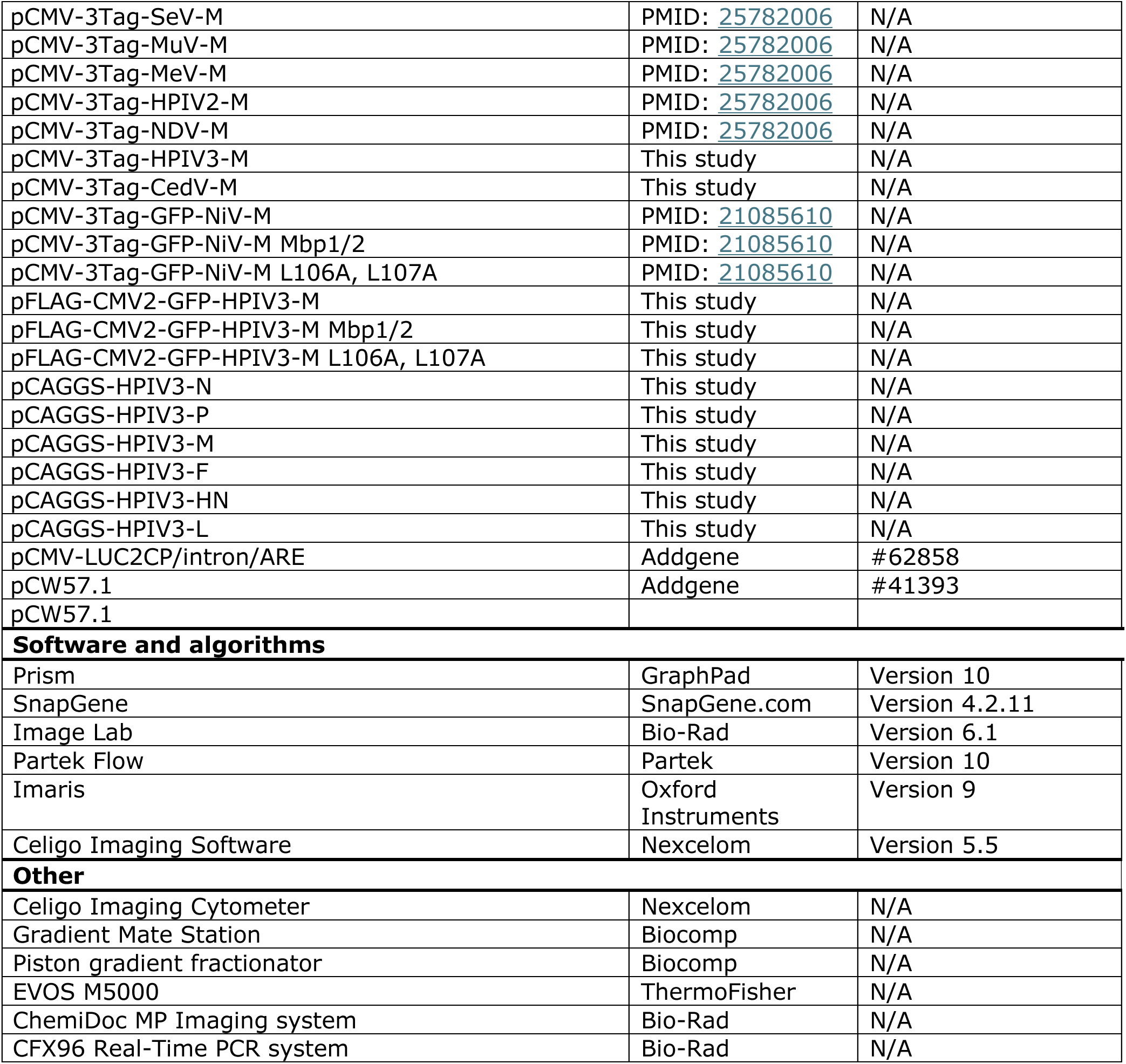

| REAGENT or RESOURCE | SOURCE | IDENTIFIER |
| --- | --- | --- |
| **Antibodies** |  |  |
| Anti-FLAG M2 antibody | Sigma-Aldrich | F3165 |
| Anti-HA antibody | Sigma-Aldrich | H3663 |
| Anti-HPIV3-N | This study | N/A |
| Anti-HPIV3-M | This study | N/A |
| Anti-eIF4AIII | Abcam | Ab180573 |
| Anti-Y14 | Abcam | Ab181038 |
| Anti-MAGOH | Abcam | Ab180505 |
| Anti-GFP | Cell Signaling | 2956 |
| Anti-S6 Ribosomal Protein | Cell Signaling | 2217, 2317 |
| Anti-L7a Ribosomal Protein | Cell Signaling | 2415 |
| Anti-GAPDH | Cell Signaling | 2118 |
| Anti-β-Actin | Cell Signaling | 3700 |
| Anti-β-Tubulin | Cell Signaling | 2128 |
| Anti-Lamin A/C | Cell Signaling | 4777 |
| Anti-COX IV | LI-COR | 926-42214 |
| Anti-Enterovirus D68 VP1 | GeneTex | GTX125989 |
| Anti-Influenza A Virus Nucleoprotein | GeneTex | GTX125989 |
| Anti-SARS-CoV2-N | Dr. Thomas Moran | Clone:1C7 |
| Goat anti-Rabbit IgG (H+L) Highly Cross-Adsorbed Secondary Antibody, Alexa Fluor 647 | ThermoFisher | A21245 |
| Goat anti-Mouse IgG (H+L) Highly Cross-Adsorbed Secondary Antibody, Alexa Fluor 647 | ThermoFisher | A21236 |
| **Bacterial and virus strains** |  |  |
| Recombinant rHPIV3-EGFP JS strain | PMID: 28405630 | GenBank: KY295925 |
| Recombinant rHPIV3-EGFP-HA-M JS strain | This study | N/A |
| Recombinant rHPIV3-EGFP-ΔM-mCherry JS strain | This study | N/A |
| Recombinant rHPIV3-EGFP-ΔM JS strain | This study | N/A |
| Recombinant rMuV-EGFP JL5 strain | PMID: 28405630 | GenBank: KY295913 |
| Recombinant rNDV-EGFP LaSota strain | PMID: 28405630 | GenBank: KY295917 |
| Recombinant rCedar-EGFP CG1a strain | This study | GenBank: JQ001776 |
| EV D68 strain US/MO/14-18947 | ATCC | VR1283 |
| Influenza A virus strain A/WSN/33 | PMID: 561860 | N/A |
| SARS-CoV-2 strain USA/WA1 | BEI resources | NR-52281 |
| Max Efficiency Stbl2 competent cells | ThermoFisher | 10268019 |
| **Chemicals, peptides, and recombinant proteins** |  |  |
| DMEM, high glucose, pyruvate | ThermoFisher | 11995-065 |
| Opti-MEM, Reduced Serum Medium | ThermoFisher | 31985-070 |
| Lipofectamine LTX and PLUS reagent | ThermoFisher | 15338100 |
| Lipofectamine RNAiMAX Transfection Reagent | ThermoFisher | 13778100 |
| BioT transfection reagent | Bioland Scientific | B01-01 |
| Fetal Bovine Serum - Optima | R&D Systems | S12450 |
| Fetal Bovine Serum - Tetracycline-free) | Takara | 631101 |
| Penicillin-Streptomycin | ThermoFisher | 15140122 |
| PBS | ThermoFisher | 10010023 |
| Trypsin-EDTA (0.25%) | ThermoFisher | 2500-056 |
| InFusion HD Cloning Kit | Takara Bio | 639650 |
| Paraformaldehyde 16% Aqueous Solution | Electron Microscopy Sciences | 15710-S |
| Intercept Blocking Buffer | Li-COR | 927-70001 |
| Collagen I-coated coverslip | Neuvitro | GG-18-15-Collagen |
| Cycloheximide (CHX) | Sigma-Aldrich | C7698-1G |
| Triton X-100 | Sigma-Aldrich | T8787 |
| Diethiothreitol (DTT) | Sigma-Aldrich | D9779 |
| Puromycin | ThermoFisher | A1113803 |
| Zeocin | ThermoFisher | R25001 |
| 4–15% Mini-PROTEAN TGX Precast Protein Gels | Bio-Rad | 4561083,4561086 |
| Trans-Blot Turbo RTA Mini 0.2 µm PVDF Transfer Kit | Bio-Rad | 1704272 |
| ProLong Glass Antifade Mountant with NucBlue Stain | ThermoFisher | P36983 |
| 1M Tris-HCl pH 8.0 | ThermoFisher | AM9855G |
| 1M Tris-HCl pH 7.5 | ThermoFisher | 15567027 |
| 0.5M EDTA pH 8.0 | ThermoFisher | 15575020 |
| 5M NaCl | ThermoFisher | AM9760G |
| 1M MgCl<sub>2</sub> | ThermoFisher | AM9530G |
| 2M KCl | ThermoFisher | AM9640G |
| RNaseOUT | ThermoFisher | 10777019 |
| Halt™ Protease Inhibitor Cocktail, EDTA-Free | ThermoFisher | 78437 |
| Trichloroacetic acid (TCA) | Sigma-Aldrich | T9159-100G |
| Anti-FLAG M2 affinity gel | Sigma-Aldrich | A2220 |
| Anti-HA Magnetic Beads | ThermoFisher | 88837 |
| **Critical commercial assays** |  |  |
| Direct-zol RNA miniprep kit | Zymo Research | R2050 |
| Luna Universal qPCR Master Mix | NEB | M3003 |
| LunaScript RT SuperMix Kit | NEB | E3010 |
| Nano-Glo HiBit lytic detection system | Promega | N3040 |
| ONE-Glo luciferase assay system | Promega | E6100 |
| **Deposited data** |  |  |
| Polysome profile mRNA sequencing data | This study | GSE274026 |
| **Experimental models: Cell lines** |  |  |
| HEK 293T | ATCC | Cat# CRL-3216 |
| HeLa | ATCC | Cat# CCL-2 |
| Vero | ATCC | Cat# CCL81 |
| Vero-E6 | ATCC | Cat# CRL-1586 |
| RD | ATCC | Cat# CCL-136 |
| Vero-HPIV3-Mpot | This study | N/A |
| BSR-T7 | PMID: 9847328 | N/A |
| BSR-T7-HPIV3-Mopt | This study | N/A |
| BSR-T7-rNiV-N-P-L replicon | This study |  |
| **Oligonucleotides** |  |  |
| See Table S1 | This study | N/A |
| **Recombinant DNA** |  |  |
| pCMV-3Tag-NiV-M | PMID: 21085610 | N/A |
| pCMV-3Tag-HeV-M | PMID: 25782006 | N/A |
| pCMV-3Tag-GhV-M | PMID: 25782006 | N/A |
| pCMV-3Tag-SeV-M | PMID: 25782006 | N/A |
| pCMV-3Tag-MuV-M | PMID: 25782006 | N/A |
| pCMV-3Tag-MeV-M | PMID: 25782006 | N/A |
| pCMV-3Tag-HPIV2-M | PMID: 25782006 | N/A |
| pCMV-3Tag-NDV-M | PMID: 25782006 | N/A |
| pCMV-3Tag-HPIV3-M | This study | N/A |
| pCMV-3Tag-CedV-M | This study | N/A |
| pCMV-3Tag-GFP-NiV-M | PMID: 21085610 | N/A |
| pCMV-3Tag-GFP-NiV-M Mbp1/2 | PMID: 21085610 | N/A |
| pCMV-3Tag-GFP-NiV-M L106A, L107A | PMID: 21085610 | N/A |
| pFLAG-CMV2-GFP-HPIV3-M | This study | N/A |
| pFLAG-CMV2-GFP-HPIV3-M Mbp1/2 | This study | N/A |
| pFLAG-CMV2-GFP-HPIV3-M L106A, L107A | This study | N/A |
| pCAGGS-HPIV3-N | This study | N/A |
| pCAGGS-HPIV3-P | This study | N/A |
| pCAGGS-HPIV3-M | This study | N/A |
| pCAGGS-HPIV3-F | This study | N/A |
| pCAGGS-HPIV3-HN | This study | N/A |
| pCAGGS-HPIV3-L | This study | N/A |
| pCMV-LUC2CP/intron/ARE | Addgene | #62858 |
| pCW57.1 | Addgene | #41393 |
| pCW57.1 |  |  |
| Software and algorithms |  |  |
| Prism | GraphPad | Version 10 |
| SnapGene | SnapGene.com | Version 4.2.11 |
| Image Lab | Bio-Rad | Version 6.1 |
| Partek Flow | Partek | Version 10 |
| Imaris | Oxford Instruments | Version 9 |
| Celigo Imaging Software | Nexcelom | Version 5.5 |
| Other |  |  |
| Celigo Imaging Cytometer | Nexcelom | N/A |
| Gradient Mate Station | Biocomp | N/A |
| Piston gradient fractionator | Biocomp | N/A |
| EVOS M5000 | ThermoFisher | N/A |
| ChemiDoc MP Imaging system | Bio-Rad | N/A |
| CFX96 Real-Time PCR system | Bio-Rad | N/A |

### RESOURCE AVAILABILITY

#### Lead Contact

Further information and requests for resources and reagents should be directed to and will be fulfilled by the lead contact, Dr. Benhur Lee

#### Materials Availability

Reagents are available from the lead contact at request.

#### Data and Code Availability

- The raw and analyzed for RNA-sequencing is accessible at NCBI GEO under the accession number GSE274026.
- This paper does not report original code.

### EXPERIMENTAL MODEL AND SUBJECT DETAILS

#### Cell lines

HEK 293T (Human embryonic kidney), HeLa cells (Human cervical carcinoma), RD cells (Human Rhabdomyosarcoma), Vero cells (African Green monkey kidney), Vero-E6 cells (Clone E6 from African Green monkey kidney) and BSR-T7 cells (a derivative of BHK-21, Syrian golden hamster kidney) were all maintained at 37°C in a 5% CO_2_ in Dulbecco’s modified Eagle’s medium (DMEM) supplemented with 10% fetal bovine serum (FBS) and 1% 100X penicillin/streptomycin solution (ThermoFisher). For confocal microscopy imaging, HeLa cells were seeded on an 18 mm #1.5 collagen I-coated coverslip (Neuvitro). For plasmid transfection, cells were transfected using Lipofectamine LTX per the manufacturer’s instructions (ThermoFisher). To generate cells with doxycycline-inducible expression of HPIV3 matrix, cDNA encoding codon optimized HPIV3-M was inserted into pCW57.1 (Addgene, #41393) through *NheI* and *AgeI* sites. Lentivirus encoding HPIV3-M was produced by co-transfecting HEK293T cells with pCMV-VSV-G, psPAX2, and pCW57.1-HPIV3-M. At 48 hours post-transfection, the supernatants were collected and centrifuged at 800 x g to remove cell debris. BSR-T7 and Vero cells were transduced with pCW57.1-HPIV3-M lentivirus. At 48 hours post-transduction, the cells were selected with 10 µg/mL puromycin (ThermoFisher, A1113803) in DMEM contain 10% tetracycline-free FBS.

#### Viruses

Recombinant viruses contain HPIV3 JS strain (GenBank: KY295925), Cedar CG1a (GenBank: JQ001776), MuV JL5 strain (GenBank: KY295913), and NDV LaSota strain (GenBank: KY295917) were propagated in Vero cells maintained in DMEM medium supplemented with 10% FBS. EV D68 strain US/MO/14-18947 (ATCC, VR-1823) were amplified in RD cells. Influenza A virus strain A/WSN/33 (ATCC, VR-1520) was propagated in embryonic eggs. SARS-CoV-2 strain USA/WA1 (BEI resources, NR-52281) was amplified in Vero-E6 cells.

### METHOD DETAILS

#### Virus inoculation and titration

For infection, cells were inoculated at the designated multiplicity of infection (m.o.i.) in serum-free DMEM. The virus was allowed to adsorb at 37°C for 1 hour, after which cells were washed with phosphate-buffered saline (PBS) and incubated at 37°C in a medium containing 10% FBS. At specific time points post-infection, the supernatant was harvested for titration, and cells were washed with PBS for cell lysate or RNA extractions. Titrations of HPVI3, Cedar, MuV, and NDV stocks were performed on Vero cells in a 96-well format, with individual infection events (infectious units, IU/mL) identified by GFP fluorescence at 24 hours post-infection using a Celigo Imaging Cytometer (Nexcelom). Plaque assay is utilized for the titration of EV D68, Influenza A, and SARS-CoV-2.

#### Plasmids and reverse genetic constructs

Cloning of codon-optimized 3X-Flag-tagged Nipah virus matrix (NiV-M), Hendra virus (HeV-M), Ghana virus (GhV-M), Sendai virus (SeV-M), Mumps virus (MuV-M), Measles virus (MeV-M), Human Parainfluenza Viruses 2 (HPIV2-M) and Newcastle disease virus M (NDV-M) is described in ^14,16^. We similarly codon optimized and cloned the open reading frames encoding M from Human Parainfluenza Viruses 3 (HPIV3-M) and Cedar virus (CedV-M), fragments were inserted within the *HindIII* and X*hoI* sites of pCMV-3Tag-1. For constructing Flag-GFP-tagged HPIV3-M, EGFP was fused to the N-terminus of M by overlapping PCR, and the cDNAs were then in-frame inserted at the *EcoRI* and *EcoRV* sites of pFLAG-CMV2. Alignment of HPIV3-M sequences using Clustal Omega identified sequence motifs corresponding to NiV-M’s nuclear export sequence (NES) and bipartite nuclear localization sequence (BpNLS) ^16^. Mutations were generated using overlap extension PCR. For Flag-tagged HPIV3 viral proteins, cDNA from HPIV3 nucleocapsid (N), phosphate (P), matrix (M), fusion protein (F), receptor binding protein (HN) and polymerase (L) were PCR fused with Flag-tag at either N- (M and P) or C-terminus (N, F, HN and L), the fragments were inserted at the *EcoRI* and *NheI* sites of pCAGGS. We modified the rHPIV3-JS construct to generate the rHPIV3-HA-M virus by inserting a HA tag at the N-terminus of the matrix protein. Additionally, we constructed the rHPIV3ΔM-mCherry and rHPIV3ΔM viruses by introducing mCherry cDNA between the P and F genes to replace the coding sequence of M or by deleting the coding sequence of M altogether.

#### Recovery of rHPIV3ΔM-mCherry and rHPIV3ΔM viruses from cDNA

For virus rescue, 4 × 10^5^ doxycycline-inducible BSR-T7 cells expressing HPIV3-M per well were seeded into a 6-well format and induced to express HPIV3-M with 500 ng/mL doxycycline The following day, transfection reactions were performed as previously described ^67^ with the plasmid encoding the antigenomic sequence of rHPIV3ΔM-mCherry or rHPIV3ΔM. The recovery of viruses was monitored (Figure S8) using EVOS M5000 imaging system (ThermoFisher); recombinant rHPIV3ΔM-mCherry and rHPIV3ΔM viruses were then amplified in doxycycline-inducible Vero cells expressing HPIV3-M.

#### Cloning of a rNiV N-P-L Replicon

The rNiV N-P-L replicon was constructed using a combination of PCR, overlap extension PCR, and InFusion cloning, based on our previously described full-length rNiV_Mal_ GLuc-P2A-eGFP reverse genetics plasmid^68^. First, the rNiV_Mal_ GLuc-P2A-eGFP construct was digested with *MluI-HF* and *AgeI-HF* overnight, followed by gel purification of the vector. InFusion cloning was then employed to re-integrate the removed NiV sequence and to incorporate a codon-optimized HiBiT-tagged eGFP reporter gene (Twist Biosciences). Next, this intermediate construct was digested with *PacI-HF* and *BsiWI-HF* restriction enzymes. Using overlap extension PCR and InFusion cloning, we restored the NiV-P gene and inserted a PuroR-P2A-BleoR gene in place of the NiV-M gene. Downstream of the integrated PuroR-P2A-BleoR gene, we maintained the NiV-M 3’ UTR up to the NiV-M gene end signal and intergenic ‘CTT’ motif. Immediately following the ‘CTT’ intergenic sequence, we appended the 5’ UTR of the NiV-L gene, starting from the NiV-L gene start signal, and restored the sequence of NiV-L through the *BsiWI* restriction site. This cloning strategy ensured that each encoded viral gene retained its native 3’ and 5’ UTRs.

#### Generating rNiV N-P-L replicon stable cells

To generate rNiV N-P-L replicon stable cells, we followed an adapted protocol as previously described ^67^. 3.8 × 10^5^ BSR-T7 cells per well were seeded into a 6-well format. The following day, cells were transfected with the plasmid encoding the antigenomic sequence of the N-P-L replicon. Cells were monitored daily for GFP-positive signals. At 72 hrs post-transfection, the cells were trypsinized and transferred to a T75 flask containing 5.0 µg/mL of puromycin (ThermoFisher, A1113803). This selection pressure was maintained until most GFP-negative cells had died. After 5 days, the medium was replaced with 150 µg/mL of zeocin, and the cells were further cultured in zeocin (ThermoFisher, R25001) until GFP-positive colonies appeared. The bulk GFP-positive population was then passaged in the presence of Zeocin and/or puromycin to ensure the stability and selection of the replicon-containing cells.

#### Deriving rNiV N-P-L replicon VLPs

To generate rNiV N-P-L replicon viral-like particles (VLPs), 3.8 × 10^5^ cells of the BSRT7 cells containing the rNiV N-P-L replicon were seeded into a 6-well format. The following day, the cells were transfected using a 1:1:1 ratio of AU1-tagged NiV-F, HA-tagged NiV-G, and untagged NiV-M plasmids. Transfection complexes were prepared by diluting 0.67 µg each of NiV-F, NiV-G, and NiV-M in 200 µL of DMEM, followed by the addition of 3 µL of BioT transfection reagent (Bioland Scientific, B01-01). After a 10 min incubation at room temperature, the transfection complexes were added dropwise to the cells. The cells were monitored for syncytia formation, with media changes every 48 hr. At 5 days post-transfection, when most of the monolayer had fused, the supernatant was collected and clarified by centrifugation at 800 x g for 10 min. The clarified supernatant was then aliquoted and frozen at -80°C until further use.

#### siRNA depletion of host factors

For siRNA depletions, cells were treated with siRNAs against eIF4AIII, Y14, and MAGOH (FlexiTube siRNA, QIAGEN) or non-targeting siRNA (Sigma-Aldrich) in a pool format (mixture of 3 siRNA targeting a single gene) for 48 h. Lipofectamine RNAiMAX (ThermoFisher, 13778100) (3.4 µL), Opti-mem (200 µL), and siRNA pool (2 µL) were mixed and incubated for 20 min at room temperature, then reverse transfections were performed in 12 well plates with 4 x 10^5^ HEK 293T cells per well at a final concentration of 20 nM siRNA.

#### Immunoprecipitation and immunoblotting

For Flag or HA tag protein immunoprecipitation, cells were harvested using lysis buffer (50 mM Tris–HCl, pH 7.4, with 150 mM NaCl, 1 mM EDTA, and 1% Triton-X-100), samples were placed on ice for 30 min, centrifuged at 12 000 × g for 30 min. Fixed amounts of cell lysate were subsequently incubated with Anti-FLAG M2 affinity gel (Sigma-Aldrich, A2220) or Anti-HA Magnetic Beads (ThermoFisher, 88837) overnight at 4°C. The reactants were washed five times with wash buffer (50 mM Tris–HCl, pH 7.4, with 150 mM NaCl) and the immunoprecipitation complex was eluted by 2× sample buffer (125 mM Tris–HCl, pH 6.8, with 4% SDS, 20% (v/v) glycerol, and 0.004% bromophenol blue at 90°C for 5 min. The eluate proteins were subjected to immunoblot analysis. All protein samples were run under reduced conditions in 1x sample buffer containing 100 mM dithiothreitol (DTT, Sigma-Aldrich, D9779). Samples were incubated in a heating block at 95°C for 10 min, resolved in a 4 to 15% SDS-PAGE gel (Bio-Rad, 4561083), and transferred to polyvinylidene difluoride (PVDF) membranes (Bio-Rad, 1704272). Membranes were blocked with phosphate-buffered saline blocking buffer (LI-COR; 927-700001) and then probed with the indicated antibodies. Antibodies against FLAG (Sigma-Aldrich, F3165), HA (Sigma-Aldrich, H3663), EGFP (Cell signaling, 2956), HPIV3-N (Benhur Lee), HPIV3-M (Benhur Lee), eIF4Alll (Abcam, ab180573), Y14 (Abcam, 2956), MAGOH (Abcam, ab180505), S6 (Cell signaling, 2217 and 2317), L7a (Cell signaling, 2415), GAPDH (Cell signaling, 2118), Actin (Cell signaling, 3700), Beta-tubulin (Cell signaling, 2128), Lamin A/C (Cell signaling, 4777), COX IV (LI-COR, 926-42214), EV D68 VP1(GeneTex, GTX132313), SARS-CoV2-N (1C7 from Thomas Moran) and Influenza A virus Nucleoprotein (GeneTex, GTX125989) were used. Membranes were washed and probed with Alexa Fluor 647-conjugated anti-mouse or anti-rabbit (ThermoFisher, A21245 and A21236). The signal of Alexa Fluor 647 was detected using the ChemiDoc MP imaging system (Bio-Rad). Relative puromycylated protein abundance was calculated by first normalizing abundance relative to Actin expression and then normalization to either mock infection or empty vector. In cytoplasmic-nuclear fractionation, relative cEJC protein abundance in each fraction was calculated by first normalizing abundance relative to the expression of the fraction marker Lamin A/C or beta-tubulin and then calculated the cytoplasm to nucleus ratios.

#### Immunofluorescence microscopy and image analysis

Cells were washed with PBS and fixed with 4% formaldehyde for 20 min at room temperature. Fixed cells were permeabilized in a blocking buffer containing PBS, 0.5% Triton X-100, and 1% BSA. After incubation with antibodies/probes in blocking buffer, samples were washed in blocking buffer and mounted on glass slides with ProLong glass antifade mountant with NucBlue stain (ThermoFisher, P36983). The slides were imaged on a Zeiss LSM 880 confocal microscope, acquiring (or without) optical Z-stacks of 0.3–0.5 µm steps. HPIV3-M was detected with rabbit anti-HPIV3-M antibodies (1:200), and cEJC was detected with rabbit anti-eIF4Alll, Y14, and MAGOH antibodies (1:500). Alexa-fluor conjugated Anti-IgG antibodies of appropriate species reactivity and fluorescence spectra were used for secondary detection (1:1000) (ThermoFisher). To determine the quantity of matrix, image analysis was performed with Imaris from Oxford instrument using the multicomponent detection module, cytoplasm and nucleus mean intensities for matrix were acquired. The statistical analysis involved calculating the ratio of the mean cytoplasmic region intensity to the mean nuclear region intensity for each cell. Given that the cytoplasmic/nuclear fluorescent intensity (C: N) ratio for the wild-type (WT) matrix is close to 1, C: N ratios greater than 1 imply increased cytoplasmic retention whereas C: N ratios less than 1 indicate increased nuclear retention. Between 30-50 cells were counted for each condition.

#### Polysome profiling

For polysome profiling, a 10-cm dish of HEK-293Ts was mock-infected or infected at a multiplicity of infection of 5 with HPIV3 for 48 hrs. Cells were treated with 100 µg/mL cycloheximide (CHX, Sigma-Aldrich, C7698) for 5 min at 37°C, then washed with cold PBS containing 100 µg/mL CHX. Cells were scraped into a 15 mL centrifuge tube and pelleted at 300 × g for 10 min. Cells were resuspended in 1 mL polysome lysis buffer (20 mM Tris-HCl pH 8.0, 100 mM KCl, 5 mM MgCl2, 1% (v/v) Triton-X100, 100 µg/ml CHX, 1mM DTT, 2U/ µL RNaseOUT and 1X EDTA-free protease inhibitor), vortexed briefly, and incubated on ice for 15 min. Later, cells were subjected to 5–7 passages through a 26-gauge syringe followed by centrifugation at 4°C for 10,000 x g for 20 min, and the clarified lysates were used for gradient sedimentation analysis. Sucrose gradient was prepared via a Gradient Mate Station (Biocomp) using 10% and 50% sucrose dissolved in polysome buffer (20 mM Tris-HCl pH 8.0, 100 mM KCl, 5 mM MgCl2, and 1X EDTA-free protease inhibitor). 30 OD of lysate was resolved on a 10–50% (wt/vol) sucrose gradient by centrifugation at 40,000 rpm at 4 °C for 150 min in a Beckman SW41-Ti rotor. 600 µL fractions were collected from the top of the gradient while monitoring absorbance at λ = 254 nm on a piston gradient fractionator (Biocomp). Total RNA was extracted from cells using a Direct-zol RNA miniprep kit (Zymo Research) and protein was trichloroacetic acid (TCA) precipitated and analyzed by immunoblotting.

#### RNA extraction, RNA-Seq, and gene expression analysis

Total RNA was extracted polysome and monosome fractions using a Direct-zol RNA miniprep kit (Zymo Research) according to the manufacturer’s protocol. Polyadenylated RNA enrichment, RNA-seq library preparation, and sequencing process were conducted at Azenta Life Sciences (South Plainfield, NJ, USA). Sequencing libraries were sequenced on an Illumina HiSeq platform (2x150bp, ∼350M pair-end reads). Gene expression analysis was performed on Partek Flow (Partek), reads were trimmed and mapped to the hg38 and rHPIV3-JS genomes, and Transcripts per Kilobase Million (TPM) were calculated for genes with mapped reads in all the fractions of both uninfected and infected cells using the total number of mapped exons reads. Density analysis was performed in python using kernel density estimation.

#### Reverse transcription and real-time quantitative PCR (RT-qPCR)

Total RNA was extracted using the Direct-zol RNA Miniprep Kit (Zymo Research, R2050). Equivalent amounts of total RNA were reverse transcribed using either oligo(dT) primers or viral genome-specific primers with the LunaScript RT Master Mix Kit (NEB, E3010).

Quantitative PCR (qPCR) was performed with gene-specific primers (Table S1) and Luna Universal qPCR Master Mix (NEB, M3003) on the Bio-Rad CFX96 Real-Time PCR system (Bio-Rad). The relative RNA levels of specific targets were normalized to GAPDH or 18S rRNA and calculated using the comparative threshold cycle (ΔΔCT) method.

### QUANTIFICATION AND STATISTICAL ANALYSIS

One-way and two-way ANOVA were used to estimate statistical significance among multiple groups and conditions, while an unpaired t-test was applied for comparisons between two groups. Data are presented as mean ± standard deviation (SD) with biological triplicates. A P-value of ≤0.05 was considered statistically significant, with significance levels indicated as follows: * P≤0.05; ** P≤0.01; *** P≤0.001; **** P≤0.0001; NS, not significant. Statistical analyses were performed using Prism 10 software (GraphPad Software).

## SUPPLEMENTAL INFORMATION TITLES AND LEGENDS

### Supplemental information

Document S1. Figures S1-S8 and Table S1

Data S1. Excel file containing data too large to fit in a PDF, related to Figure 3

Data S2. Excel file containing data too large to fit in a PDF, related to Figure 4

**Figure S1.**
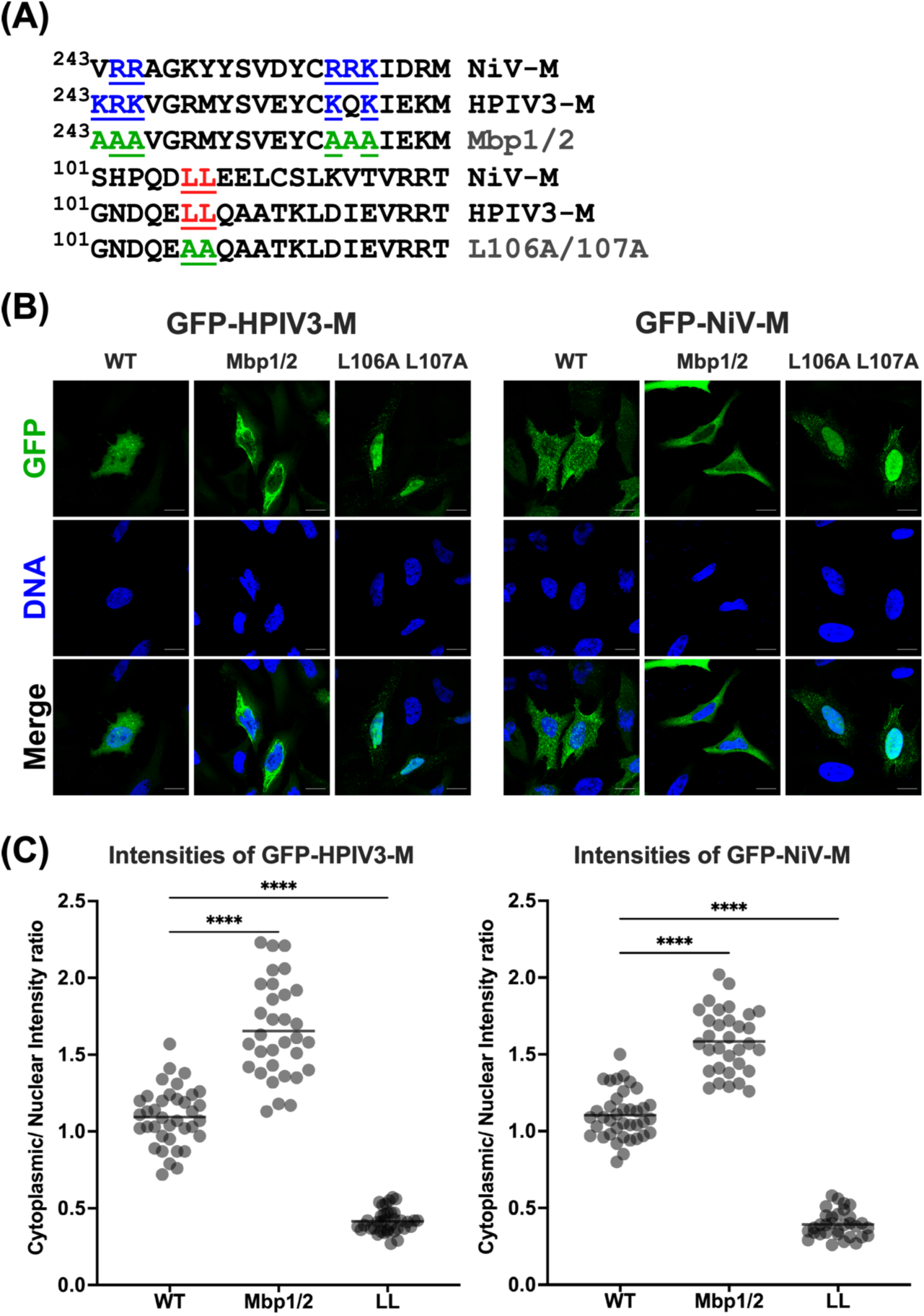
Mutagenesis studies of nuclear localization signals (NLSs) and nuclear export signals (NESs) in GFP fused HPIV3-M and NiV-M. (A) Positively charged amino acid residues in the bipartite NLSs or key leucine residues in the potential NESs were mutated to alanine. (B) HeLa cells expressing either wild-type (WT), NLS mutant (Mbp1/2), or NES mutant (L106A L107A) forms of GFP-fused HPIV3-M and NiV-M were fixed and stained with Hoechst to visualize nuclei. Representative fields of cells expressing each construct are shown. Scale bars represent 20 µm (C) Quantification of the cytoplasmic/nuclear GFP intensity (C: N) ratios for 30–50 individual cells was analyzed for each mutant, as described in the Materials and Methods. Statistical significance was determined by one-way ANOVA with Dunnett’s multiple comparison test. **** P < 0.0001.

**Figure S2.**
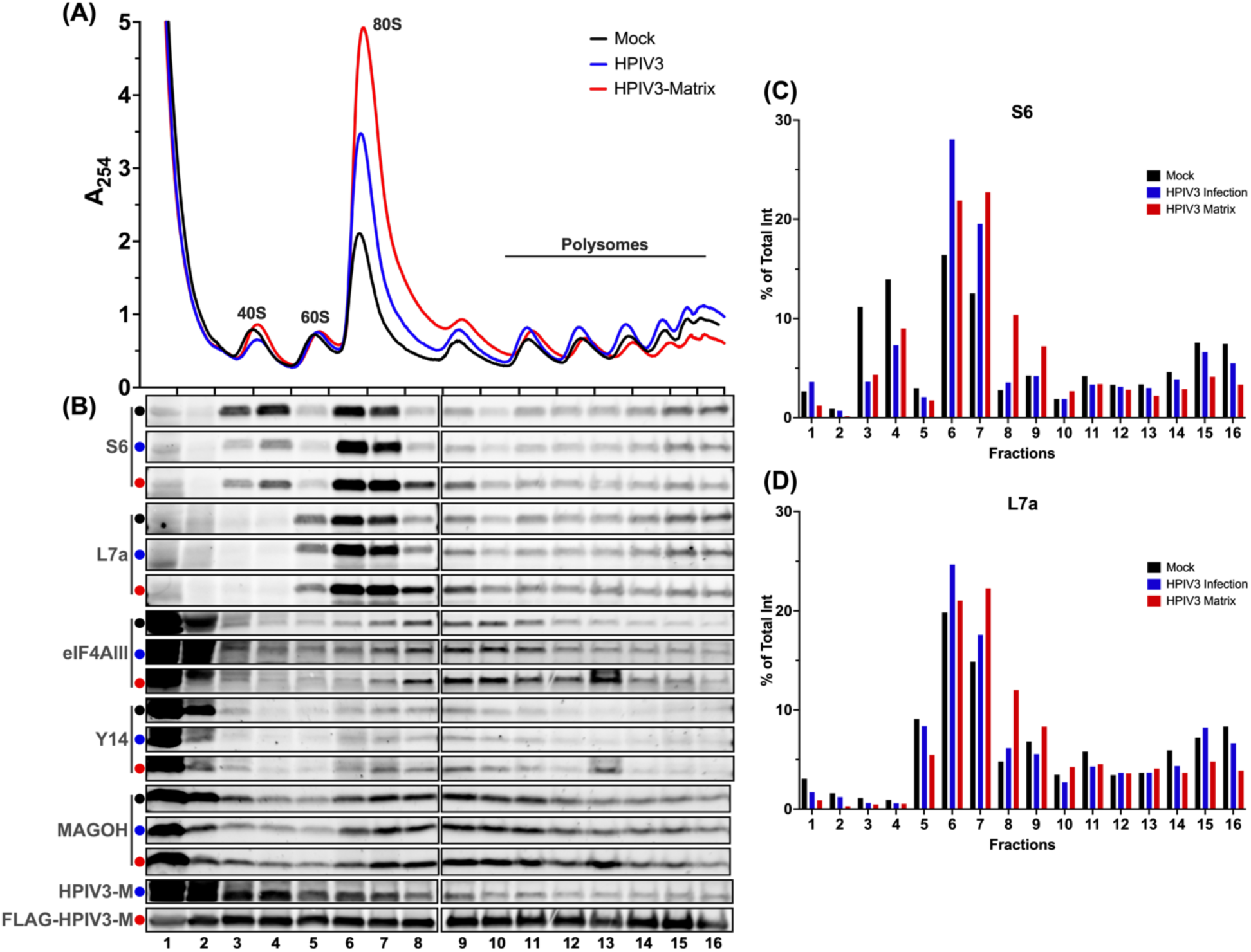
Effects of HPIV3 infection and matrix protein on host translational profile. (A) Polysome profiles of mock-infected (black), HPIV3 infected (blue), or HPIV3-M transfected (red) HEK-293Ts at 48 hrs post-infection. HEK-293Ts were infected with HPIV3 (MOI of 5) or transfected with matrix for 48 hrs and cytoplasmic extracts were prepared for polysome profiling. Cytoplasmic extracts were sedimented through a 10–50% sucrose gradient and 0.6 ml fractions were collected while continuously measuring absorbance at λ = 254nm. (B) Proteins were TCA precipitated from the collected fraction with equal volume and analyzed by immunoblotting to determine the sedimentation of S6, L7a, eIF4Alll, Y14, MAGOH, and HPIV3-M with ribosomal subunits, monosomes, or polysomes. (C-D) Densitometric quantification of the indicated proteins (S6 and L7a) across 16 fractions from (B). The y-axis shows the percentage of the total integrated intensity (% of Total Int) for each protein in the indicated condition: mock infection (black), HPIV3 infection (blue), and HPIV3 matrix protein expression (red).

**Figure S3.**
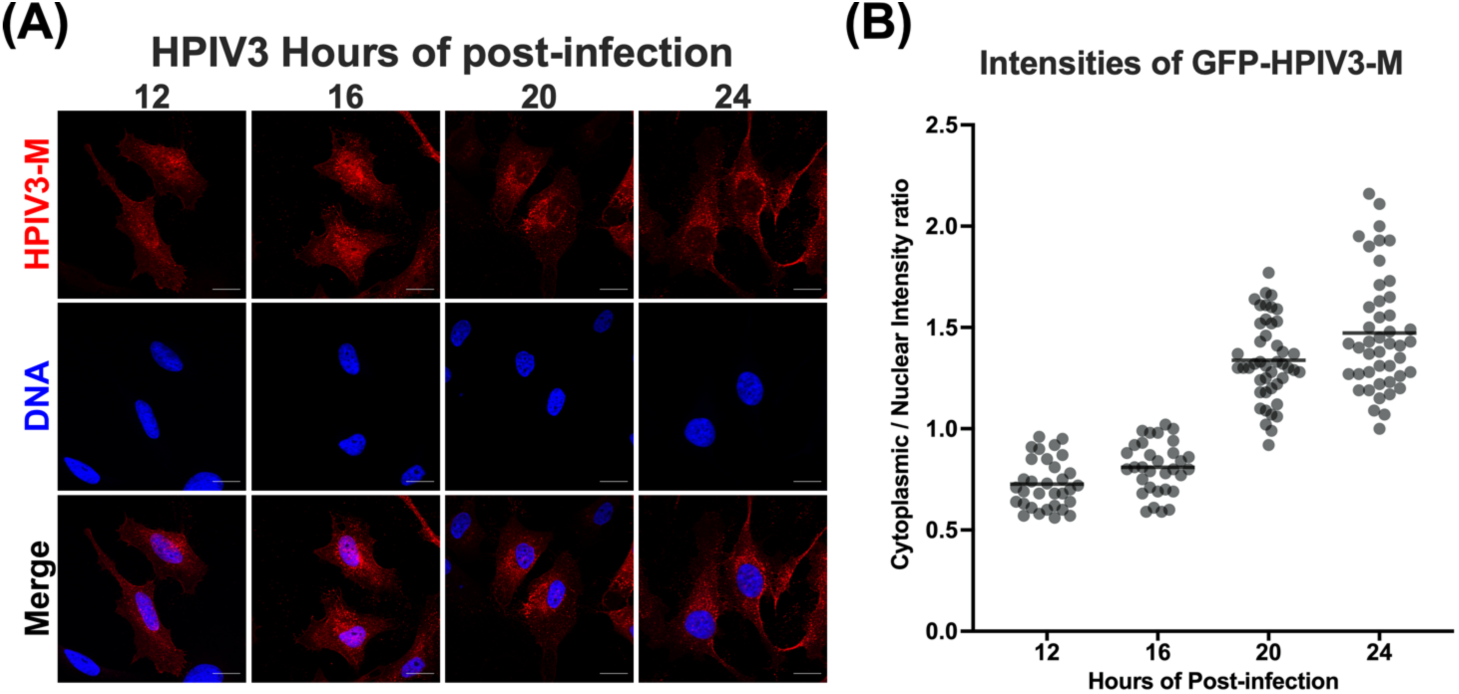
Nuclear-cytoplasmic trafficking of HPIV3 matrix protein (HPIV3-M) during infection. (A) HeLa cells infected with HPIV3 at m.o.i of 5 and then incubated with fresh growth medium for up to 24 hrs. At 12, 16, 20, and 24 hours of post-infection, cells were fixed and counterstained with anti-HPIV3-M antibody (red) to label viral matrix protein, and nuclei were stained with Hoechst (blue). Representative fields of cells at each time point are shown. Scale bars represent 20 µm (B) Quantification of cytoplasmic/nuclear HPIV3-M intensity (C: N) ratios was performed on 30–50 individual cells, as described in the Materials and Methods. Statistical significance was analyzed by one-way ANOVA with Dunnett. **** P <0.0001; ns, not significant.

**Figure S4.**
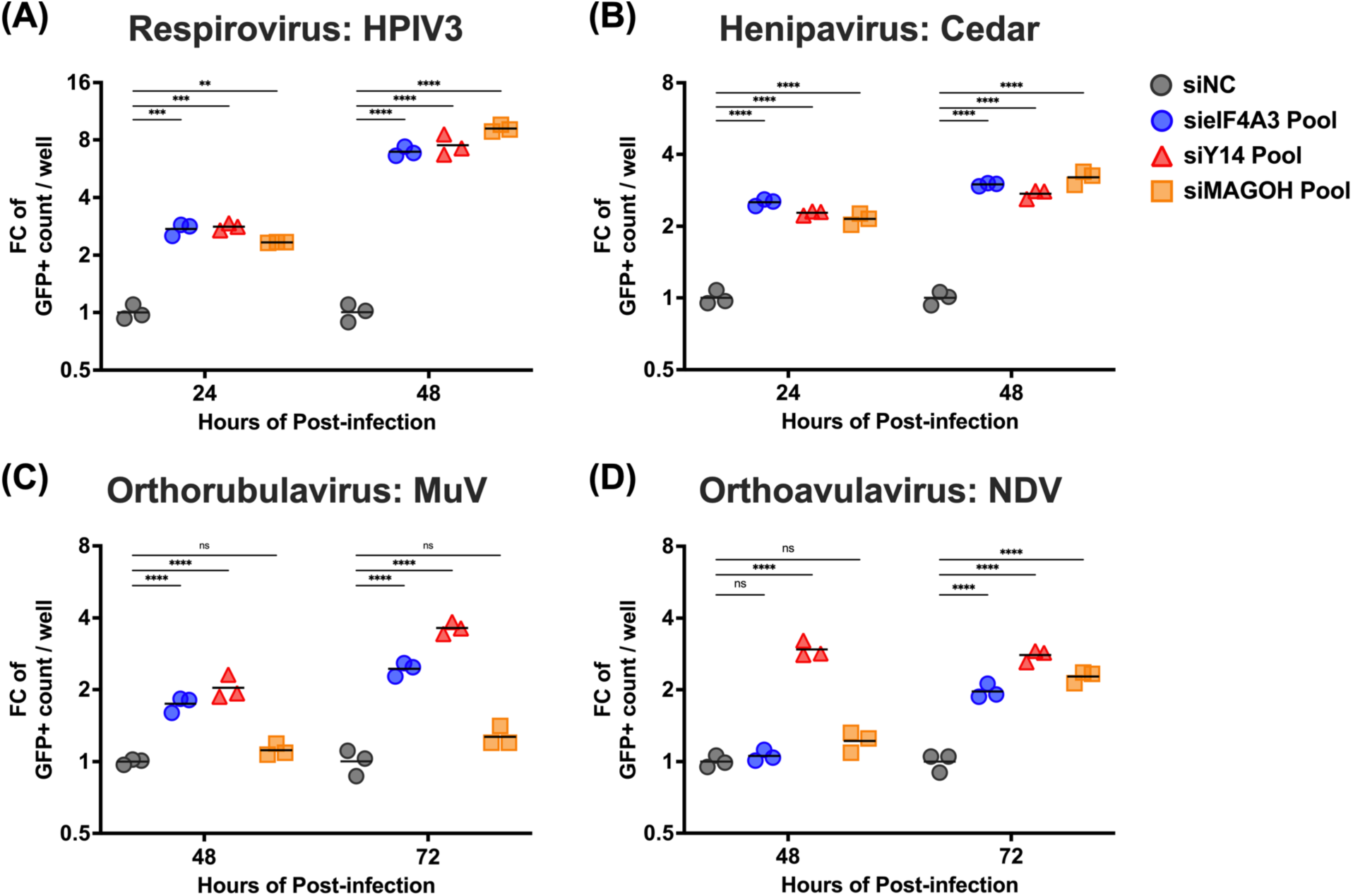
Effects of the exon junction complex on Paramyxovirus infection. HEK-293T cells were transfected with siRNA pools targeting eIF4A3, Y14, MAGOH, or non-targeting control siRNAs (NC), respectively. At 48 hrs post-transfection, cells were inoculated with the designated virus (A) HPIV3, (B) Cedar, (C) MuV, and (D) NDV at a multiplicity of infection (m.o.i.) of 0.01. The number of GFP-positive cells in each well was acquired at the indicated time points by Celigo imaging cytometer (Nexcelom). Relative fold changes (FC) in GFP-positive cells per well were then calculated. Symbols represent the data points from biological triplicates. Bars represent the mean of the triplicates. Statistical significance was determined by two-way ANOVA with Dunnett multiple comparison test. * *P* <0.05; ** *P* <0.01; *** *P* <0.001; **** *P* <0.0001; ns, not significant.

**Figure S5.**
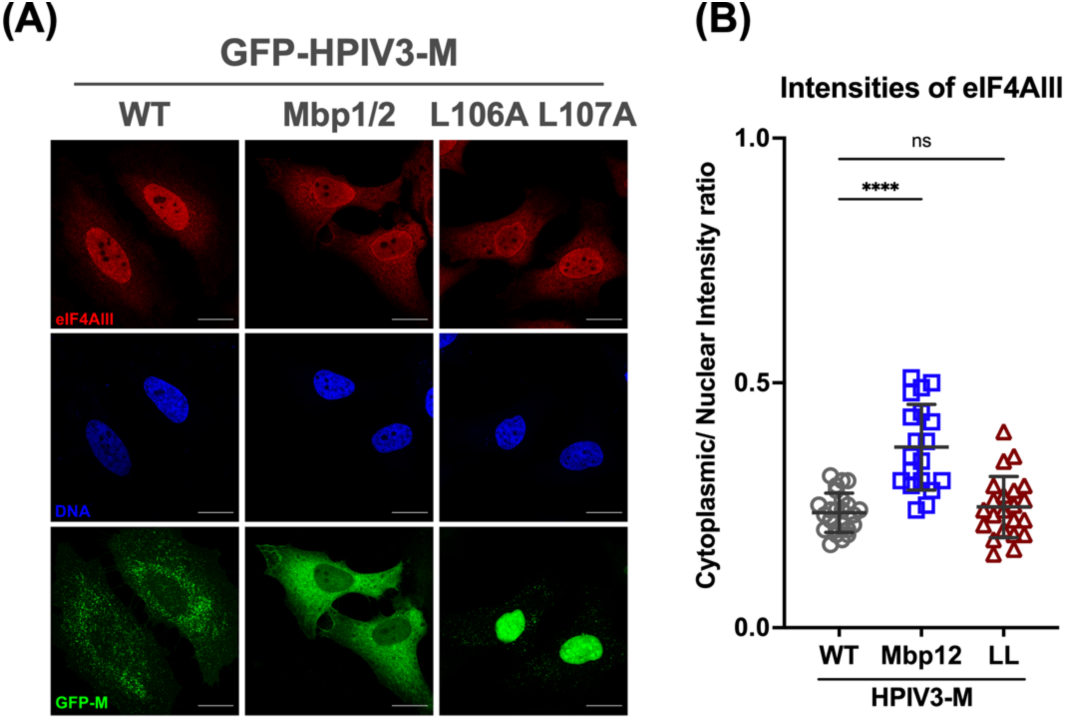
Subcellular localization of eIF4Alll in HeLa cells expressing GFP fused HPIV3 matrixes. Left panel HeLa cells expressing either wild-type (WT), NLS mutant (Nbp1/2), or NES mutant (L106A L107A) forms of GFP-fused HPIV3-M and NiV-M were fixed and stained with anti-eIF4AIII (red) and Hoechst for nuclei (blue), and GFP fluorescence indicates M expression. Representative fields of cells for each condition are shown. Scale bars represent 20 µm. Right panel: Quantification of cytoplasmic/nuclear eIF4Alll intensity (C: N) ratios was performed on 30 individual cells, as described in Materials and Methods. Statistical significance was analyzed by one-way ANOVA with Dunnett multiple comparison test. **** *P* <0.0001; ns, not significant.

**Figure S6.**
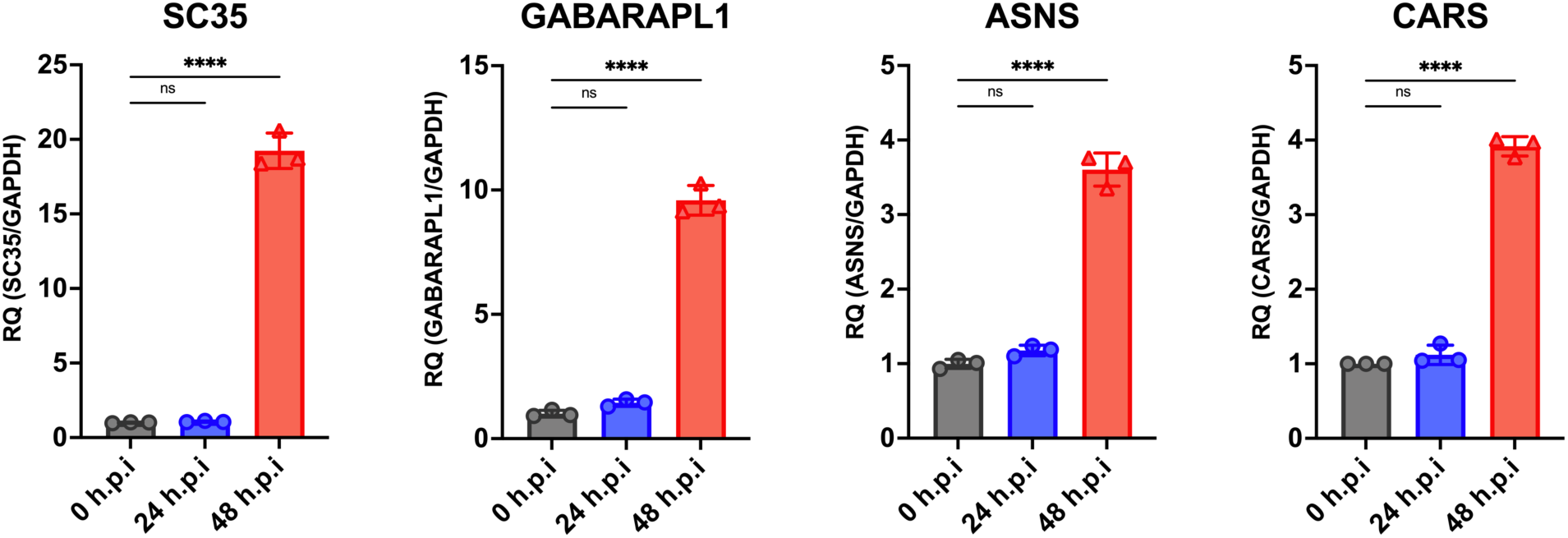
Effects of HPIV3 infection on NMD activity. HEK-293T cells infected with HPIV3 at m.o.i of 0.01 and analyzed at 24 and 48 hours of post-infection. Endogenous targets of the NMD mRNA surveillance pathway, SC35, GABARAPL1, ASNS, and CARS were analyzed by quantitative RT-PCR. Relative quantification (Gene/GAPDH) is normalized to uninfected controls. Symbols are data points from biological triplicates. Bar represents the mean ± SD. Statistical significance was determined by one-way ANOVA with Dunnett multiple comparison test. * P <0.05; **** P <0.0001; ns, not significant.

**Figure S7.**
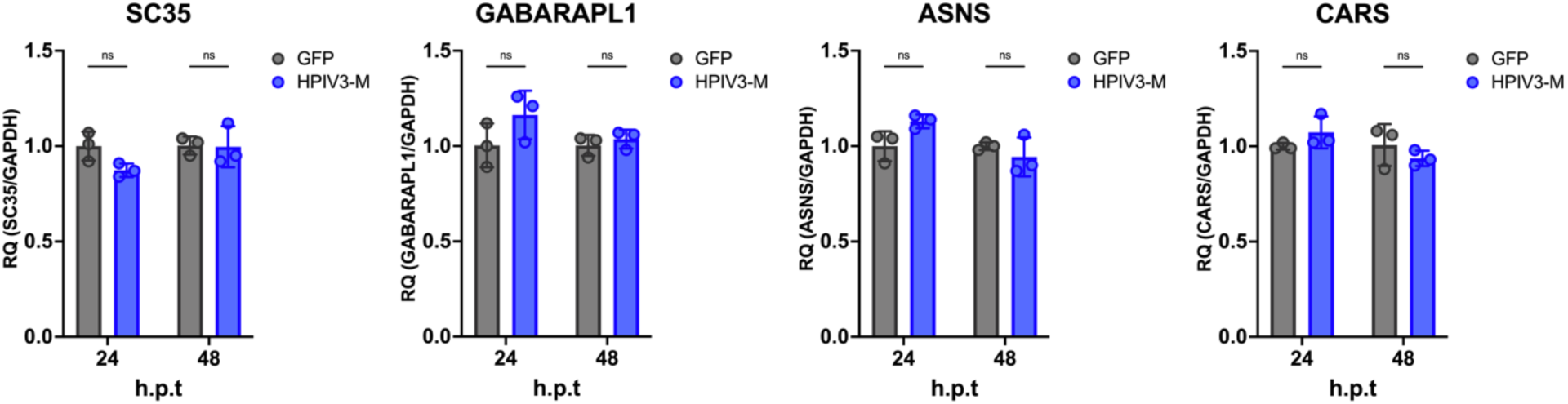
Effects of HPIV3 matrix on NMD activity. HEK-293T cells were transfected with the indicated protein and analyzed at 24 and 48 hrs post-transfection. Endogenous targets of the NMD mRNA surveillance pathway, SC35, GABARAPL1, ASNS, and CARS were analyzed by quantitative RT-PCR. Relative quantification (Gene/GAPDH) is normalized to uninfected controls. Symbols are data points from biological triplicates. Bar represents the mean ± SD. Statistical significance was determined by two-way ANOVA with Bonferroni’s multiple comparisons test. ns, not significant.

**Figure S8.**
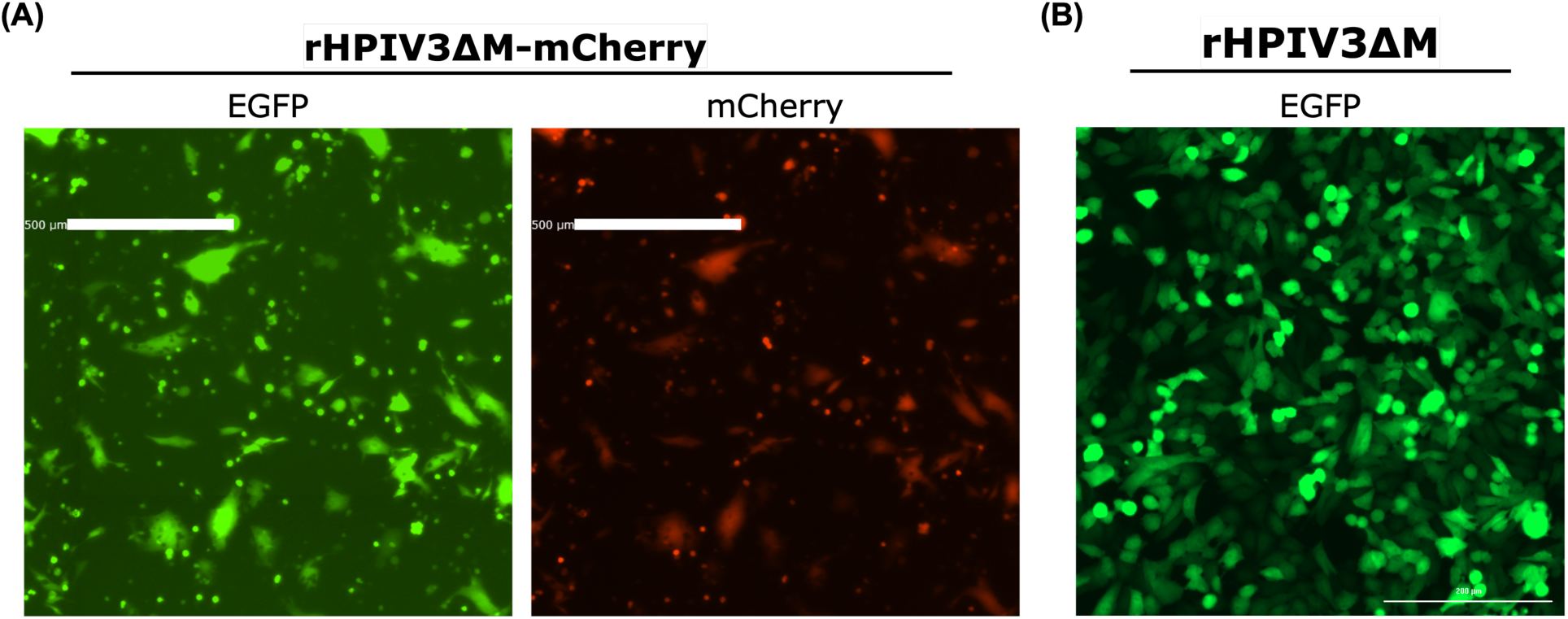
Recovery of rHPIV3ΔM-mCherry and rHPIV3ΔM viruses in BSR-T7 cells. Representative images from the rescue of (A) rHPIV3ΔM-mCherry and (B) rHPIV3ΔM at day 3 of post-transfection in BSR-T7 cells. Images were captured by EVOS m5000.

**Table S1:**
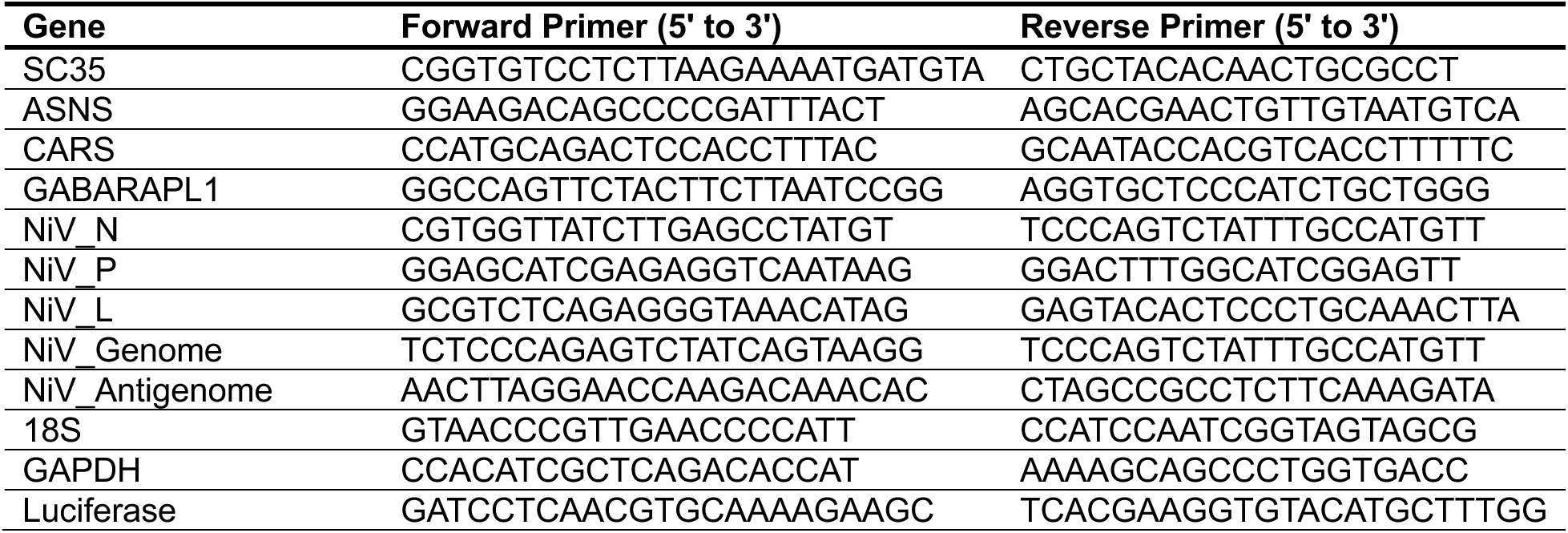
qPCR primers, related to Star Methods.

**Data S1: Mapped viral read counts of polysome profile mRNA sequencing for HPIV3-WT and HPIV3-delta-M virus.**

Tab 1: TPM normalized viral genes expression for HPIV3-WT and HPIV3-delta-M virus.

Tab 2: Ribosome association efficiency of viral transcripts for HPIV3-WT and HPIV3-delta-M virus.

**Data S2. Mapped read count of polysome profile mRNA sequencing for Mock, HPIV3-WT and HPIV3-delta-M samples.**

